# Structural basis for activation and potentiation in a human α5β3 GABA_A_ receptor

**DOI:** 10.1101/2025.01.27.635004

**Authors:** John Cowgill, Chen Fan, Jan Stayaert, Rebecca J. Howard, Erik Lindahl

## Abstract

General anesthetics like etomidate mediate amnesic as well as sedative effects through different populations of gamma amino butyric acid receptors (GABA_A_Rs) in the central nervous system. The amnesic effects have largely been attributed to α5-subunit-containing receptors in the hippocampus, indicating a clear role in learning and memory for these receptors. The α5 subunit is thought to primarily coassemble with β3 and, in some cases, γ2 subunits, generating a variety of receptor subtypes with differential functional and pharmacological properties. However, the stoichiometry, structure, and gating mechanisms of these different subpopulations are not well understood. Here we report structures of human α5β3 GABA_A_Rs with various modulators, assembled in two stoichiometries. Our cryo-EM structures, combined with electrophysiology in *Xenopus* oocytes, support a primary assembly of 2:3 α:β subunits, though a minority population of 1:4 α:β indicates multiple assemblies are possible. Differential glycosylation of the α5 and β3 subunits enabled reconstruction of the heteromeric complex even in the absence of added fiducials. In the resting state of the receptor, Zn^2+^ binds to histidine residues at the M2-17′ position in the β3 subunit, blocking ion passage. In the activated state, GABA binding to the orthosteric site is associated with global rearrangements propagating to unbinding of the 17′ Zn^2+^ atoms and opening of the 9′ hydrophobic gate. Unlike the α1β3 receptor where GABA is effectively a partial agonist, saturating GABA binding to α5β3 appears to drive activation of nearly all receptors, resulting in a single desensitized state observed under the cryo-EM conditions. In agreement with this high GABA efficacy, the GABA-bound structure is virtually unaffected by further addition of the anesthetic etomidate at the β-α transmembrane domain interface. These structures offer new detailed models for gating and modulation of a GABA_A_R subtype critical to learning and memory, including prospective templates for structure-based drug discovery.

## INTRODUCTION

Type-A γ-aminobutyric acid receptors (GABA_A_Rs) are pentameric ligand-gated ion channels that drive the majority of the inhibitory neurotransmission in the brain by passing a hyperpolarizing chloride current in response to binding GABA. These receptors control a diverse range of physiological processes through both phasic and tonic inhibition mediated by distinct populations of synaptic and extrasynaptic receptors, respectively^1,2^. GABA_A_Rs are critical drug targets as exemplified by their role as the primary targets of general anesthetics like etomidate^3,4^. Though therapeutic doses of etomidate induce loss of consciousness, subtherapeutic doses potently block memory formation without inducing general anesthesia^5–7^. Subtype-specific inhibitors and genetic knockouts have attributed this amnesic effect of etomidate to GABA_A_ receptors containing the α5 subunit^5–7^. Thus there has been considerable interest in designing α5-specific modulators for conditions associated with learning and memory deficits like Down syndrome, Alzheimers, depression, and schizophrenia^8^.

There are 19 genes encoding GABA_A_Rs subunits in humans (α1–6, β1–3, γ1–3, δ, ε, ρ1–3, π and θ) that each correspond to an N-terminal extracellular domain (ECD) and a transmembrane domain (TMD) of four transmembrane helices (M1–M4)^9^. The ion conduction pathway is formed at the central axis of the receptor and is lined by the second transmembrane helix (M2) of each of the five subunits in the receptor. The neurotransmitter-binding site is formed at the subunit interface in the ECD specifically between β and α subunits, therefore α5 must coassemble with β subunits to form functional receptors. Coexpression patterns and electrophysiology in recombinant expression systems indicate α5 coassembles with β3 and, in some cases, γ2 subunits, generating a variety of receptor subtypes with differential functional and pharmacological properties^10–13^. For example, α5β3 receptors are susceptible to zinc block while α5β3γ2 receptors are sensitive to numerous α5-specific benzodiazepines^11^.

There is substantial interest in understanding the structural determinants of gating and pharmacology of α5-containing GABA_A_Rs given the therapeutic potential of these receptors. Several studies have probed binding sites for modulators including neurosteroids and benzodiazepines in chimeric or homomeric α5 receptors engineered to improve structural tractability^14,15^. These structures have revealed critical insights into the binding modes and molecular determinants of α5-specificity in potential therapeutics, but the mechanisms of channel gating and potentiation remained unexplored due to a lack of GABA binding site in these receptors. The extrasynaptic α5β3 GABA_A_Rs was previously resolved in the unexpected stoichiometry of 1:4 α5:β3 ratio in a single GABA-bound, putatively open state^16^. More recently, the related α1β3 GABA_A_Rs was determined in the expected 2:3 α1:β3 stoichiometry in nanodiscs in the presence of GABA and/or various inhibitors^17^. Interestingly, the pore remained in a resting-like conformation even when bound to GABA. Thus there are numerous gaps in our understanding of the assembly stoichiometry and mechanisms of gating and potentiation of αβ-type GABA_A_Rs, especially those containing the α5 subunit, and whether these receptors can exhibit diversity in stoichiometry or always assemble in specific combinations.

Here, we combined cryogenic electron microscopy (cryo-EM) in both detergent micelles and lipid nanodiscs with functional analysis through electrophysiology to investigate open questions of αβ-type GABA_A_R stoichiometry, gating, and potentiation by etomidate. We find that the predominant stoichiometry in our hands is 2:3 α5:β3, though a minority class in the 1:4 stoichiometry was observable in a large dataset. Unlike in α1β3 receptors, GABA binding drives these channels into a desensitized state with high efficacy under saturating conditions. Etomidate binds in lipid-facing cavities formed during channel activation specifically in the TMD at β-α interfaces, associated with expansion of the TMD including opening of the activation gates, consistent with a conformational selection mechanism of potentiation.

## RESULTS

### Structures of human α5β3 GABA_A_Rs in resting and desensitized states

To gain insights into the stoichiometry of assembly and mechanisms of gating and potentiation of the human α5β3 GABA_A_Rs, we determined cryo-EM structures of heteromeric receptors in both detergent micelles and brain-lipid/saposin nanodiscs (Fig. 1, Extended Data Fig 1). In accordance with previous studies on αβ receptors^16,17^, and to improve biochemical tractability, the intracellular domains (M3-M4 loops) of α5 and β3 were removed and affinity tags were inserted at the N- or C-termini respectively (Extended Data Fig. 2). Furthermore, superfolder GFP was added to the N-terminus of α5 while mKalama was inserted into the M3-M4 loop of β3 to enable tracking of each subunit by fluorescence-detected size exclusion chromatography (FSEC), producing a construct hereafter called α5β3-EM. The vestibular glycosylation of α subunits prevents assembly of homomeric α5 receptors (Extended Data Fig. 3a), thus we used a single affinity purification step targeting the TwinStrep tag of α5. The affinity-purified product from HEK cells coexpressing the modified α5 and β3 subunits elute as a monodisperse peak in SEC in either detergent micelles or lipid nanodiscs, and contain both α5 and β3 as indicated by the coelution of sfGFP and mKalama in FSEC (Extended Data Fig. 3b-d).

**Figure 1.**
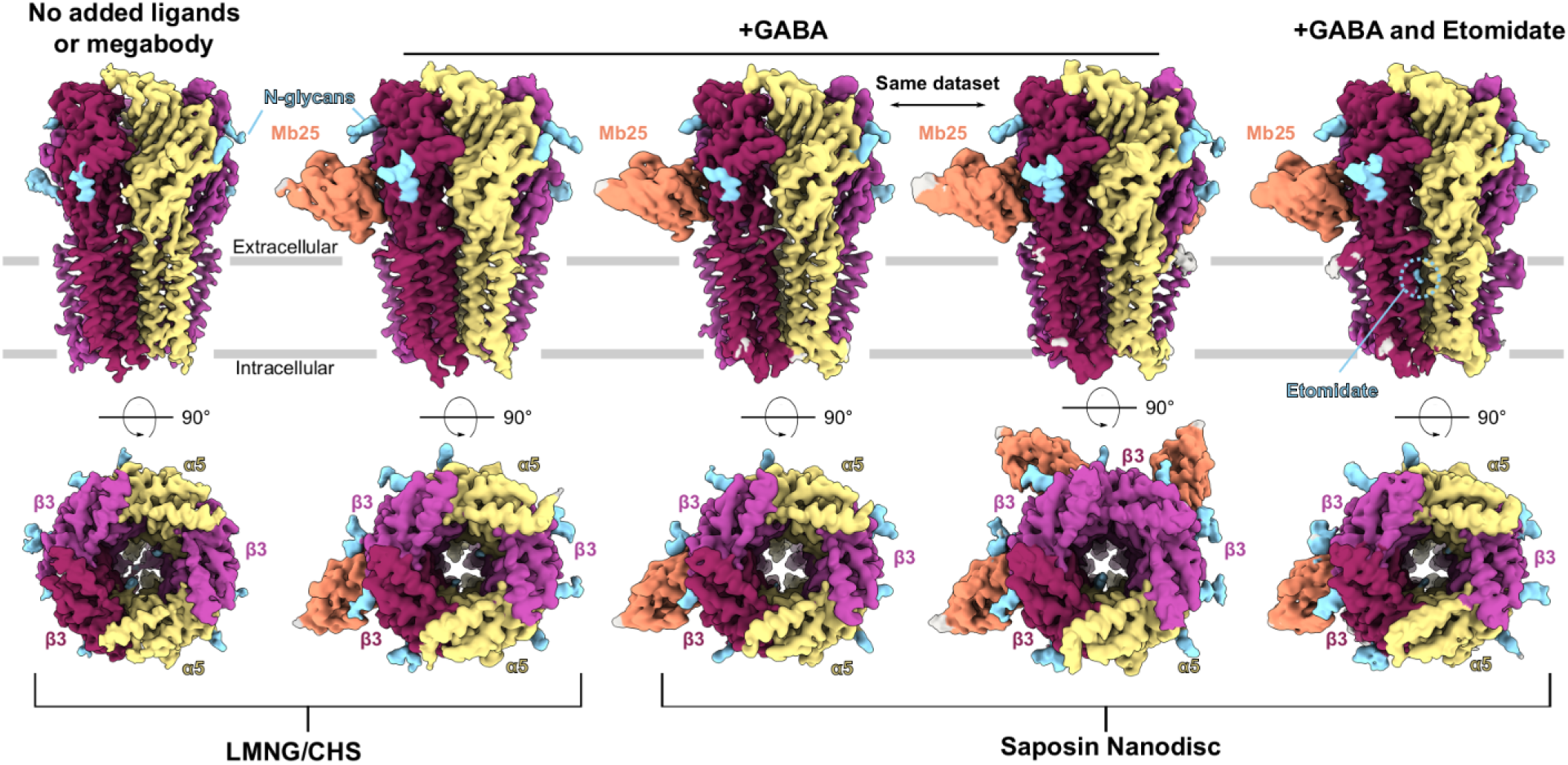
Structures of human α5β3 GABA_A_Rs in resting and desensitized states. Cryo-EM density maps for the α5β3 GABA_A_R resolved under various conditions. Maps are colored by subunit with α5 shown in yellow and β3 in light/dark magenta. Megabody 25 was used in the GABA-bound conditions and is colored in salmon. N-linked glycans and added ligands are colored in blue while unmodeled density is colored gray. Top row represents the side view of the receptors as they would sit in the membrane while the bottom row shows the top view.

Density maps from α5β3-EM showed clear distinction between α5 and β3 subunits based on the differential glycosylation pattern and binding of megabody 25 (Mb25) specifically at the β-β interface. Symmetry expansion followed by iterative focused classification on opposing ECDs enabled reconstruction of a single stoichiometry of 2:3 α5:β3 in the absence of added ligands and without addition of the Mb25 fiducial (Fig. 1, Extended Data Fig. 1b). Despite the presence of non-protein densities discussed below, this structure appeared to occupy a resting state. The dominant stoichiometry of 2:3 α5:β3 was consistent across multiple rounds of expression and purification, and confirmed in the GABA-bound datasets, where Mb25 was used to facilitate particle alignment. However, a minority class of 1:4 α5:β3 was observed in a larger dataset (47,000 micrographs) in the presence of Mb25 and a short application of GABA prior to grid freezing.

To assess the stoichiometry in wild-type α5β3 GABA_A_Rs, we measured GABA concentration-response curves in *Xenopus laevis* oocytes coexpressing α5 and β3 (Extended Data Fig. 3e). These responses were well-fitted by a Boltzmann curve with a Hill slope of 1.69 (95% confidence interval of 1.38 to 2.11). While the Hill slope is not a direct measure of the number of ligand binding sites, it is generally considered a lower bound for the number of sites present. Importantly, Hill slopes above 1 have also been reported for wild-type α5β3 GABA_A_Rs in L929 and HEK cells, strongly supporting the presence of two GABA binding sites and thereby a stoichiometry of 2:3^11,18^ for functional receptors. Similar characterization of α5β3-EM yielded a Hill slope of 1.83 (95% confidence interval of 1.49 to 2.28), indicating the high cooperativity of gating seen in the wild type is retained in our EM construct, though the apparent affinity for GABA was lower relative to the wild type (EC50 of 6.97 μM compared to 1.84 μM, respectively). We focus the majority of our structural analyses below on the receptors with 2:3 stoichiometry.

### GABA drives α5β3 into the desensitized state with high efficacy

The ion conduction pathway in the resting state of α5β3-EM is blocked by a clear density near the 17’ position of the three β-subunits, where primes denote the position on M2 relative to a conserved basic residue at the intracellular end (Extended Data Fig. 4a). This block recalls the coordination of Zn^2+^ observed in the α1β3 GABA_A_R, and our resting state largely resembles the cobratoxin-inhibited structure of that subtype^17^ (Extended Data Fig 4b-d). Though we purified the receptor in the absence of divalent cations, we assigned this density as Zn^2+^ since it is present in the expression media and is the most physiologically relevant divalent ion for these receptors. This density is not observed in our activated-state structures, which dilate substantially at this position (Fig. 2a,b).

**Figure 2.**
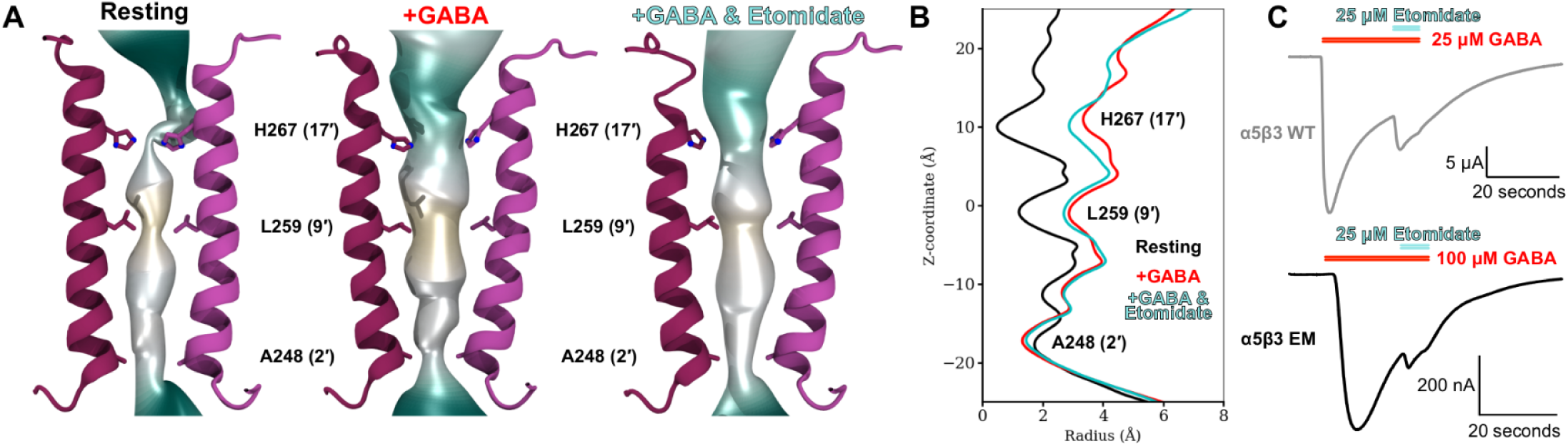
**High efficacy channel activation by GABA** (A) Structural overlay of pore pathways with opposing β subunits for the resting (left), GABA+Mb25-bound in detergent, and GABA+Etomidate+Mb25-bound structure in saposin. Pore pathway is colored by the hydrophobicity and key residues along the pore lining M2 helix are represented as sticks. (B) Pore radius as a function of Z-position relative to the 9 activation gate for the structures shown in panel A. Traces are colored with the resting state shown in black, GABA+Mb25 bound in red, and GABA+Mb25+etomidate in cyan. (C) Example traces of α5β3 WT (gray, top) and α5β3-EM (black, bottom) in response to a saturating pulse of GABA with coapplication of 25 M etomidate at the end of the pulse.

Activation of α5β3-EM by GABA results in overall expansion of the pore-lining M2 helices that is non-uniform along the ion permeation pathway (Fig. 2a,b). The dilation of the pore in the activated state is greatest in the outward-facing half of (upper) M2, while the inward-facing half of (lower) M2 becomes even more constricted than the resting state (Fig. 2b). The activation gate at the 9’ position (L271 in α5 and L259 in β3) expands to a radius of 3 Å in the GABA-bound structures (Fig. 2b). The profiles of the GABA-bound states in detergent micelles and lipid nanodiscs are largely similar, suggesting the α5β3 GABA_A_R is less sensitive to lipid mimetic than the α1βXγ2 GABA_A_Rs, which collapse under several conditions^19,20^. (Extended Data Fig. 4e,f). We also find a similar diameter of the 9’ gate in the small fraction of receptors in the 1:4 stoichiometry (Extended Data Fig. 4g,h). The pore radius at 2’ position is below 2 Å and too narrow to pass a hydrated chloride ion in all of our GABA-bound structures, consistent with a desensitized state for these receptors. The funnel-shaped ion conduction pathways in the GABA-bound structures closely resemble the profile of the α1β2γ2 GABA_A_R bound to GABA and etomidate^21^ (Extended Data Fig. 4i,j) and is characteristic of the desensitized state reported for other receptors in the pentameric ligand-gated channel family^22–26^.

A previously reported structure for α5β3 was described as an open state^16^ (Extended Data Fig. 4g,h), while the related α1β3 was best described as a primed state^17^, with a rotated ECD but the 9’ gate remaining closed (Extended Data Fig. 4k,l). To assess the physiological relevance of our GABA-bound structures, we measured the response of the wild-type and EM α5β3 constructs to saturating pulses of GABA followed by a pulse of etomidate (Fig. 2c). An initial pulse of saturating GABA leads to a transient peak in current amplitude that decays in the sustained presence of GABA. This is characteristic of channel desensitization, and indicates a large fraction of channels desensitized within the minimum 10-15 second window of GABA application prior to freezing our cryo-EM grids. Co-application of etomidate at the end of the initial GABA pulse results in an additional smaller peak in current that decays prior to washout of both GABA and etomidate (Fig. 2c). This is most likely the result of activation of a small fraction of receptors that were not driven into the open or desensitized state by GABA alone, as was previously reported for the α1β3 GABA_A_R^17^. Using the ratio of the current elicited by GABA alone to the sum of the peaks in response to GABA and etomidate, we can approximate the fraction of receptors activated by GABA alone (Extended Data Fig. 4m). This analysis shows that over 80% of the available channels are directly driven into the open and desensitized states by GABA alone (Extended Data Fig. 4n). In agreement with the limited effect of etomidate in the presence of saturating GABA, little change is observed in the pore profile in the etomidate-bound structure (Fig. 2a,b). Combined, these analyses imply the desensitized state is the expected state of the channel under the conditions of grid preparation, in agreement with the structural features we highlight above.

### Neurotransmitter binding pocket constricts and rigidifies upon GABA binding

Despite preparation of the resting state receptor in the absence of GABA, we found a serendipitous density in the neurotransmitter binding pocket, similar to those reported for α1β3γ2 and α4β3δ GABA_A_Rs^22,23^ (Extended Data Fig. 5a). As previously proposed^22^, this density could be tentatively assigned to butyrate, which is added to media during expression and only lacks the primary amine group of GABA (Figure S5). Indeed, application of butyrate opposes GABA-induced currents in a reversible manner for both the wild type and EM constructs, thus we modeled butyrate into the resting state structure (Fig. 3a, Extended Data Fig. 5a,b).

**Figure 3.**
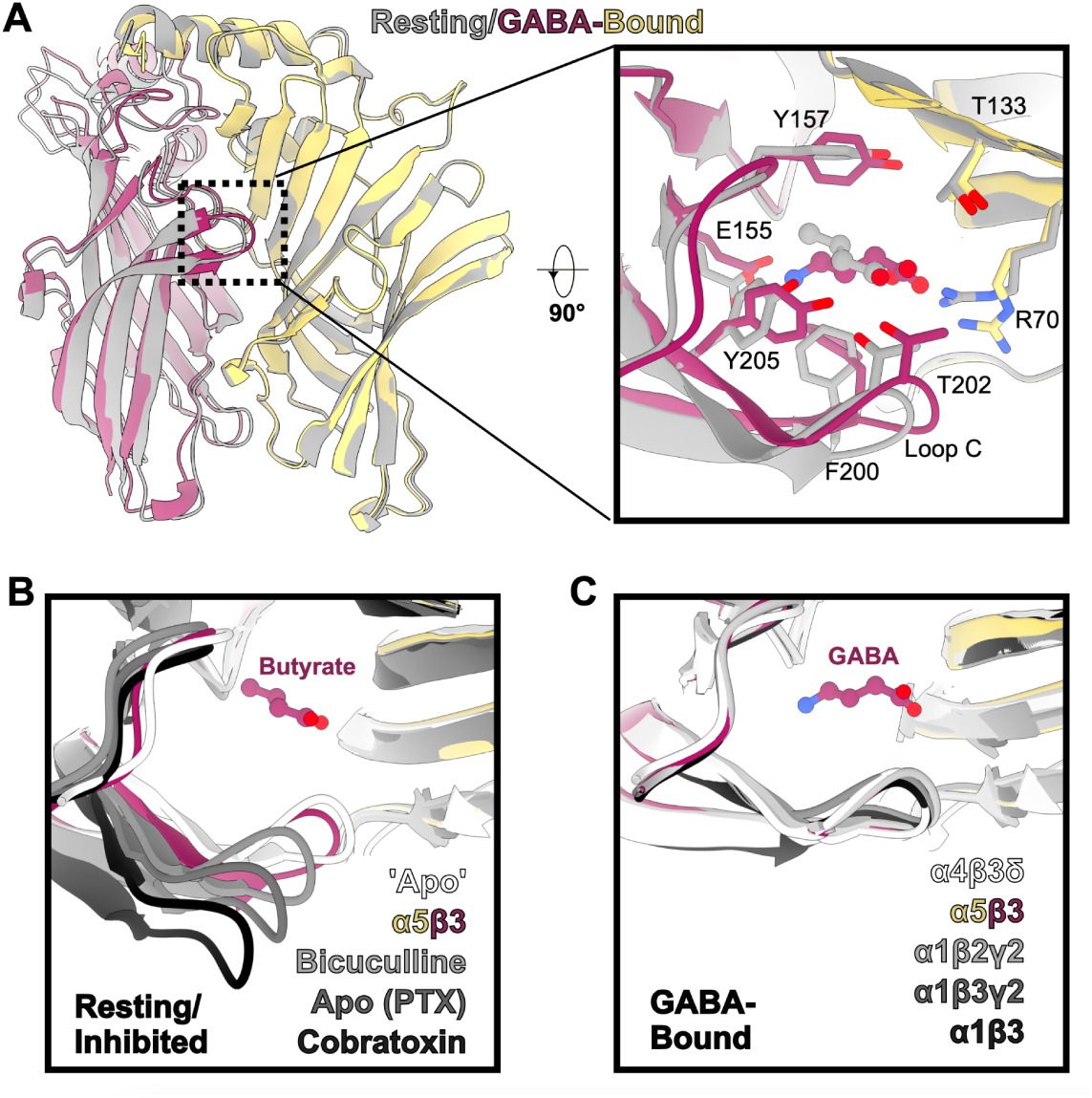
Compaction of neurotransmitter binding pocket during activation. (A) Left: ECDs of the β-α interface in the resting (gray) and desensitized (magenta/yellow) α5β3-EM structures, aligned on the complementary α5 subunits. Right: Zoomed view of the neurotransmitter binding pocket occupied by butyrate (gray) in the resting state and GABA (magenta) in the desensitized state. Relevant residues in the pocket interacting with the ligands are shown as sticks. (B) Zoomed view of the resting state of α5β3 aligned with other resting/inhibited GABA_A_R states. The receptors are colored according to the inlaid legend for the following structures: resting α4β3δ with putative butyrate density, Bicuculline-bound α1β2γ2, PTX-bound α1β3γ2, and cobratoxin-bound α1β3. Subunits are aligned based on the ECD of the complementary α subunits. (C) Zoomed view of the GABA-bound α5β3 aligned with other GABA-bound GABA_A_Rs. The receptors are colored according to the inlaid legend for the following structures: α4β3δ, α1β2γ2, α1β3γ2, and α1β3. Subunits are aligned based on the ECD of the complementary α subunits.

To visualize local changes in the neurotransmitter binding site upon GABA binding, we superimposed resting and desensitized α5β3-EM from detergent micelles, aligning by the complementary α subunit (Fig. 3a). The carboxylate groups of GABA and butyrate occupy a similar position, coordinated by T202 from loop C of the principal subunit and R69 and T132 from the complementary subunit. The aromatic box (Y157, F200, and Y205 of the principal subunit) undergoes rearrangement upon GABA binding to coordinate the primary amine of GABA together with E155. The increased electrostatic interactions with GABA compared to butyrate compress the neurotransmitter binding pocket via enhanced loop C lockdown. This is demonstrated by both a 1.4 Å shift in the tip of loop C further capping the pocket as well as reduced latent heterogeneity in density maps around the orthosteric site as shown by OccuPy^27^ (Extended Data Fig. 5d). Local scale is a normalized measure of the local degradation of contrast within a density map that depends on local heterogeneity due to structural flexibility, variable occupancy, or particle misalignment. Despite the disparity in the reconstruction resolutions, local scale values are similar in the GABA+Mb25 density maps in detergent and lipid nanodiscs. Moreover, elevated local scale values throughout the ECD in the GABA+Mb25 maps indicate reduced heterogeneity compared to the TMD region in these maps. In the resting state, local scale is dramatically reduced specifically in the ECD, consistent with increased ECD flexibility in the absence of GABA, especially in the loop C region of the neurotransmitter binding site.

Comparison of our resting α5β3-EM structure with neurotransmitter binding pockets of other GABA_A_Rs in resting or inhibited states reveals substantial heterogeneity in the extent of loop C uncapping (Fig. 3b). Our α5β3-EM is most comparable to the α4β3δ GABA_A_R which also contains a serendipitous density similar to butyrate^22^. Larger inhibitors like bicuculline^21,28^ (α1β2γ2) and cobratoxin^17^ (α1β3) are associated with increased uncapping of loop C. These and other larger competitive inhibitors are also effective against α5-containing receptors, suggesting the neurotransmitter binding pocket of α5β3 can also sample these more uncapped conformations. This is consistent with the reduced local scale in this region shown in the OccuPy analysis above. Surprisingly, the α1β3γ2 GABA_A_R bound to the non-competitive inhibitor picrotoxin (PTX) shows one of the greatest extents of uncapping despite lacking any ligands or serendipitous density in the neurotransmitter pocket^28^. Thus the apo binding pocket of α5β3 may be even further uncapped than in our resting state structure bound to butyrate. Interestingly, the β-β interface in resting α5β3-EM shows a similar uncapped conformation of loop C as the α1β3γ2 GABA_A_R structure with picrotoxin (Extended Data Fig. 6a). This interface is occupied by HEPES in both our resting and GABA (+ Mb25) bound structures of α5β3-EM, and undergoes a capping motion during receptor activation similar to the β-α interface (Fig. 3d, Extended Data Fig. 5a). However, it is unclear if this rearrangement is due primarily to binding of GABA or Mb25. Unlike the β-α and β-β interfaces, the α-β interfaces show little rearrangement upon GABA binding in our α5β3-EM structures (Extended Data Fig. 6c).

In contrast to the resting/inhibited states, the GABA-bound loop C conformation of α5β3-EM is consistent with the other heteromeric GABA_A_Rs (Fig. 3c). GABA adopts a similar pose in the binding pocket of these heteromeric receptors (Extended Data Fig. 5e), consistent with the similarity in the surrounding binding pocket and the general lack of subtype-specificity between agonists in heteromeric receptors.

### Receptor activation by GABA

GABA binding drives global movements that are generally conserved across GABA_A_R subtypes, and can be visualized by aligning the TMD regions of the resting and desensitized structures (Fig. 4). Capping of loop C over the agonist binding pocket is associated with a counter-clockwise rotation of the ECD relative to the TMD (viewed from the extracellular side) and a contraction of the upper ECD (Fig. 4A). The combined caping and rotation displaces the tip of loop C to a greater extent in β- than in α-subunits. Compared to previously reported structures of the α1β3 GABA_A_R, the GABA+Mb25-bound ECD is more rotated in our α5β3-EM structures (Extended Data Fig. 7a). The GABA-induced compaction of the ECD increases the buried surface area at all interfaces (Table 2), especially those with a principal β-subunit, in agreement with the stabilization of the ECD by GABA and Mb25 evident in the OccuPy analysis above (Extended Data Fig. 5e).

**Figure 4.**
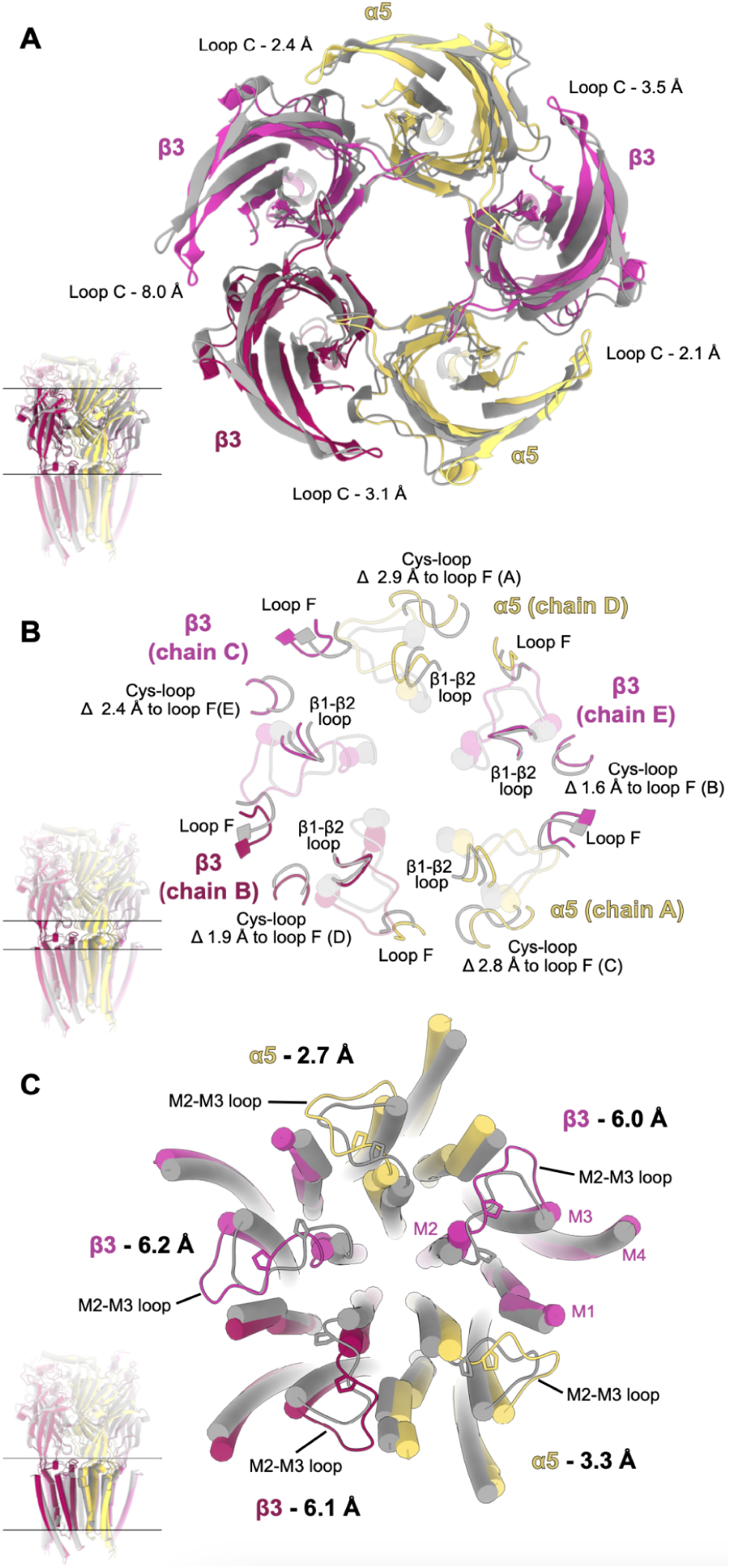
**Global rearrangements upon GABA binding** (A) Overlay of ECDs from resting (gray) and desensitized (magenta/yellow) receptors aligned by their TMDs. The slice of the receptor depicted is shown by the inset side view at the lower left. Distances measure the distances between CA atoms of the threonine at the tip of loop C. (B) Overlay of ECD-TMD interfaces from resting (gray) and desensitized (magenta/yellow) receptors aligned by their TMDs. The slice of the receptor depicted is shown by the inset side view at the lower left. Distances measure the changes in distance from the Cys-loop to loop F of subunit i+2 as defined by the CA of P150(α5)/P144(β3) on the Cys loops and Q192(α5)/Q185(β3) on loop F. (C) Overlay of TMD from resting (gray) and desensitized (magenta/yellow) receptors aligned by their TMDs. The slice of the receptor depicted is shown by the inset side view at the lower left. Distances measure the distance moved by P281(α5)/P273(β3) from the resting to desensitized state.

Counter-clockwise rotation propagates from the upper to the lower ECD, which interfaces with the M2-M3 loop of the TMD. The lower ECD also expands upon GABA binding, driving opposing subunits 1.6-2.9 Å further apart, as measured by the distance between the Cys-loop and loop F for subunits i to i+2 (Fig. 4b). This expansion of the lower ECD and contraction of the upper ECD is associated with a change in the tilt of each subunit ECD relative to the same-subunit TMD that is more pronounced in β subunits compared to α and for α5β3-EM than the α1β3 GABA_A_R (Extended Data Fig. 7c,d). Structural rearrangements in the lower ECD are most pronounced for the Cys-loop and loop F, with more minor and variable rearrangements in the β1-β2 loops. Surprisingly, the β1-β2 loops in the principal β-subunits binding GABA are associated with the least change in this region, in contrast to the α1β3 GABA_A_R where divergence is greatest in the GABA-binding β-subunits.

The rearrangements in the ECD are coupled to outward movements of the M2-M3 loop of each subunit, 0.8-2.1 Å larger than previously observed in the α1β3 GABA_A_R (Fig. 4c, Extended Data Fig. 7b). As also seen in the α1β3 subtype, the outward movement is more limited for α-subunits than for β-subunits in α5β3-EM. The M2-M3 loop displacement dilates the upper pore, and propagates down the transmembrane helices, weakening the subunit interfaces within the TMD (Table 2). The reduction in buried surface area in the TMD upon GABA binding largely offsets the increased surface burial at the ECD interfaces, so that the net change in interfacial area is only 2.3% of the total area. The decreased interfacial surface area in the TMD stems partially from the dilation of the M2 helices upon channel activation detailed above, and additionally through opening of lipid-facing cavities at the M3-M1 interfaces throughout the receptor.

The differences between α5β3 and α1β3 noted above in the state of the pore and global conformation of the receptor are surprising given the similarity in sequences of the α5 and α1 subunits (Fig. 2). These sequences are especially similar in the regions coupling the ECD to the TMD like the β1-β2 loop, Cys-loop, Loop F, and M1-M2 loops (Extended Data Fig. 8a). One notable difference is T56 in the β1-β2 loop α5 that is a histidine in α1. Interestingly, this histidine (H58) interacts with the backbone of the top of the M2 helix in both resting and GABA-bound structures of α1β3 (Extended Data Fig. 8b). In the α1β2γ2 structure bound to GABA and etomidate, channel activation is accompanied by a rotameric flip and further dilation of the top of M2 that breaks this interaction (Extended Data Fig. 8c). The smaller side chain of T56 does not reach the top of M2 even in the resting state (Extended Data Fig. 8b,c), likely reducing the barrier to channel activation.

### Etomidate stabilizes lipid-facing cavities at β-α TMD interfaces

In the presence of etomidate, we observed non-protein densities in the TMD at both of the M3-M1 interfaces between β (principal) and α (complementary) subunits that could be confidently modeled as etomidate (Fig. 5a). These densities are distinct from those observed at the α-β interfaces, or at the β-α interfaces in the absence of etomidate (Extended Data Fig. 9a-c). The resolved etomidate pose at each β-α interface of α5β3-EM resembles previously reported poses from cryo-EM structures of the α1β2γ2^21^ and homopentameric β3 GABA_A_Rs^29^, as well as poses predicted from photoaffinity labeling^30^ (Extended Data Fig. 9d-f). The binding pocket is largely hydrophobic, with M286, F289, and V290 from β3-M3 lining the principal face, and I231, L235, P236, and M239 from α5-M1 forming the complementary face. Hydrophilic residues from β3-M2 (T262 and N265) line the most buried cleft of the cavity, though these are too far to directly engage in hydrogen bonding interactions with etomidate. Interestingly, N265 has been shown to be a critical determinant of etomidate selectivity in GABA_A_Rs as mutation to a serine that is present at this position in β1 subunits eliminates potentiation by etomidate. Given the similar polar nature of serine and asparagine, the electrostatic interactions of the amide nitrogen with the phenyl ring of etomidate are likely important for binding or potentiation as previously proposed.

**Figure 5.**
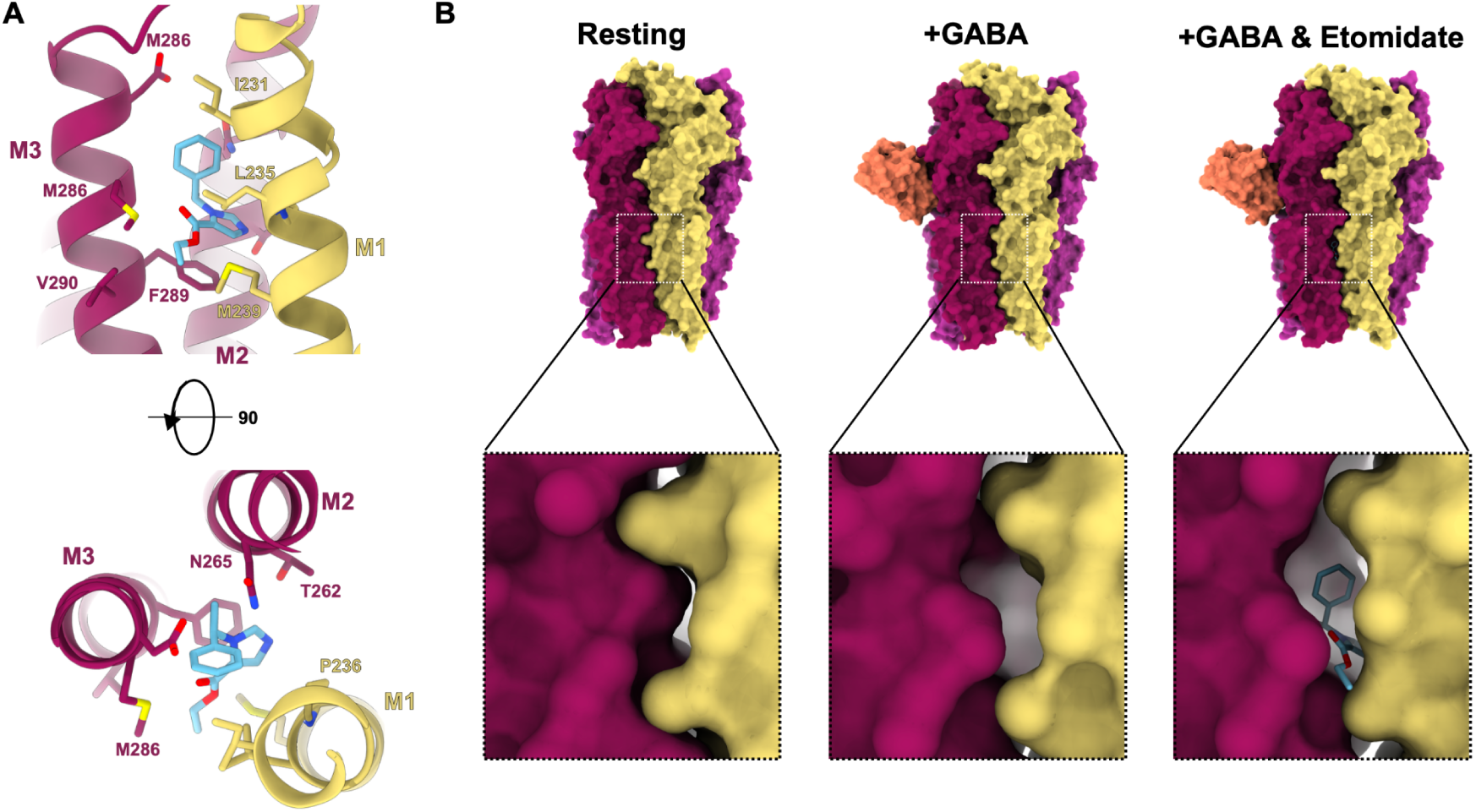
**Etomidate stabilizes lipid-facing cavities at β-α interface** (A) Zoomed view of etomidate binding site with key residues shown as sticks represented as a side view and rotated by 90° for a view from the extracellular side (bottom). (B) Surface representations of resting (left), GABA-bound (middle), and GABA+etomidate-bound (right) receptors colored by subunit. The bottom shows a zoomed in view of the subunit interface where etomidate binds in a cavity that is present in the GABA-bound structure but not in the resting state.

The etomidate binding site is occluded in the resting state by the tight packing of the β-α TMDs (Fig. 5b). The expansion of the TMD upon GABA activation opens up the etomidate binding pocket even in the presence of GABA alone (Fig. 5b). Etomidate binding to this interface causes little rearrangement at either the local or global scale (Fig. 5b,c). This is distinct from the mechanism previously observed for α1β2γ2 which showed that etomidate binding induces further rearrangement of the TMD compared to GABA alone. This is most apparent in the pore profiles above where our α5β3-EM structures are nearly identical across all of the GABA-bound structures regardless of the presence of etomidate. In contrast, α1β2γ2 shows further dilation upon etomidate binding, with the GABA-only structure intermediate between our resting and activated α5β3-EM structures (Extended Data Fig. 4j).

There is also weak density at the β-β interface that could stem from etomidate that is either more dynamic or bound at lower occupancy than at the β-α interface, though it could not be confidently built (Extended Data Fig. 9c). The TMD of the principal β subunit at the β-β interface is poorly resolved relative to other subunits, suggesting the absence of clear etomidate density here could stem from heterogeneity in the surrounding pocket. Interestingly, a similar reduced quality of density in the TMD was previously reported for the γ2 subunit in structures of the α1β2γ2 GABA_A_R^21^, which occupies the same position as the principal subunit of the β-β interface.

## DISCUSSION

Our structural and functional characterization above supports a mechanistic model for the activation and potentiation of extrasynaptic α5β3 GABA_A_Rs. In the resting state, the pore is prone to Zn^2+^ block due to the close proximity of three histidines at the top of the M2 helices in β subunits, similar to the α1β3 GABA_A_R^17^ (Fig. 6a). Loop C is more dynamic and uncapped over the neurotransmitter pocket at rest, making the receptor susceptible to inhibitors of a wide range in size, such as butyrate and bicuculline. GABA binding in the β-α interface stabilizes loop C and induces capping of the neurotransmitter pocket that stabilizes the subunit interface in the ECD (Fig. 6b). Compaction of the ECD is coupled with high efficacy to expansion of the TMD interfaces, specifically in the M2 pore lining helices and between M3 and M1 at all interfaces. The dilation of M2 opens activation gates in the middle of M2 enabling passage of hydrated ions. However, the lower gate in our structures remains closed, consistent with the desensitized state expected from macroscopic recordings. Expansions at the M3-M1 subunit interface open lipid-facing cavities in all subunits, but etomidate preferentially binds in the β-α interfaces to stabilize the activated form of the TMD (Fig. 6c). Together, these results highlight a gating mechanism that is generally conserved with other GABA_A_Rs characterized to date^21,22,28^, but also provide insights into receptor assembly and subtype-specific features of α5β3 GABA_A_R gating.

**Figure 6.**
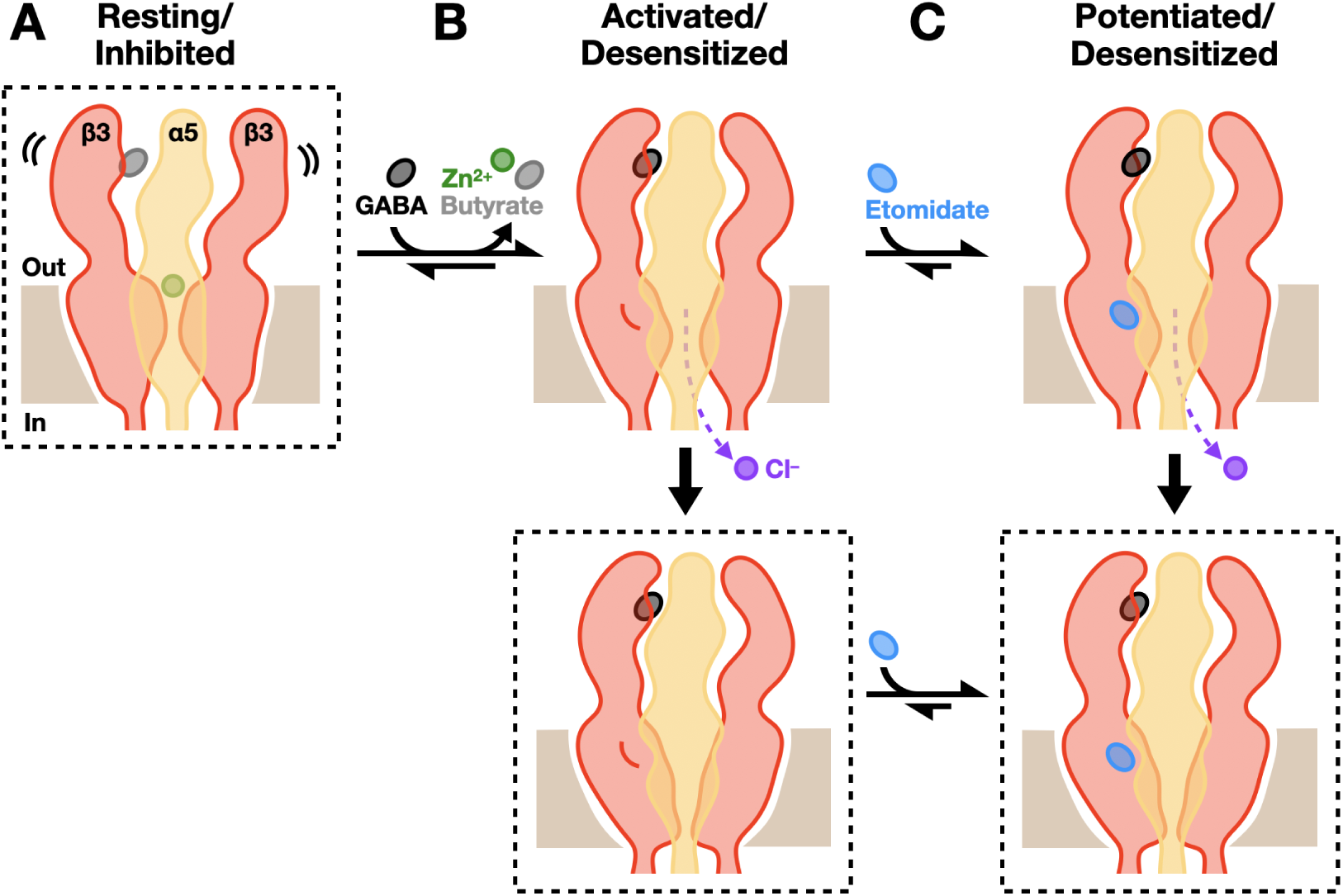
**Mechanistic model of gating and allosteric modulation in ɑ5-containing GABA_A_Rs.** (A) Schematic of a GABA_A_R containing ɑ5 (yellow) and β3 (red) subunits in a resting-like state, embedded in a plasma membrane (tan); the distal two subunits are hidden for clarity. This state features a relatively expanded, flexible ECD, and is capable of coordinating Zn^2+^ ions (green) in the outer transmembrane pore, as well as the inhibitor butyrate (gray) in the orthosteric ligand site at the extracellular β-ɑ interface. (B) Schematic as in *A* of activated structures induced by exposure to agonist, showing a presumably transient open state capable of conducting Cl^−^ ions (purple) (top), and a longer-lived desensitized state evident in cryo-EM preparations (bottom). GABA (black) displaces butyrate in the orthosteric site, and stimulates conformational transitions that relatively contract, rotate, and rigidify the ECD, dissociate Zn^2+^, and expand the transmembrane pore. (C) Schematic as in *B* of potentiated complexes in which the general anesthetic etomidate (blue) relatively stabilizes the open (top) or desensitized (bottom) state by binding a state-dependent pocket at the transmembrane β-ɑ interface. In *A–C*, dashed boxes indicate structures reported in this work.

Our cryo-EM structures, and the high cooperativity evident in GABA responses, support a predominant stoichiometry of 2:3 α5:β3 subunits. This result is consistent with the stoichiometry obtained for α1β3^17^, though it contrasts with previous work showing only a single stoichiometry of 1:4 for α5β3^16^. Both of these studies used dual affinity purification, with α5β3 purified on α and β subunits, while the α1β3 GABA_A_R was purified based on unique tags on two separate α subunit constructs. We used a single affinity purification on the basis of α subunits, as they do not form homopentamers, to minimize loss of receptors during purification and other biases to the observed stoichiometry. We do find a minor fraction of receptors in the 1:4 stoichiometry, demonstrating that both assemblies are possible for the α5β3 GABA_A_R. As past studies have suggested that GABA_A_Rs form a variety of assemblies to diversify the functional and physiological properties *in vivo*, our results may provide notable documentation of a single receptor subtype assembling with multiple stoichiometries^22,31^. Still, given that our receptors were produced in a recombinant heterologous overexpression system, factors such as relative expression levels, association with scaffolding proteins, and folding chaperones may further influence the relative abundance of the 2:3 and 1:4 assemblies in native cells.

In our GABA-bound α5β3 GABA_A_R structures, we obtain only a single conformation corresponding to the desensitized state of the receptor. Unlike the αβγ, the global structure is preserved in detergent micelles and lipid nanodiscs, consistent with the single channel recordings of αβ-type GABA_A_Rs have shown that the open state in these receptors is very short-lived, with open durations of less than 10 ms. Previous structural work on α1β3 showed that GABA activates the channel with low efficacy. Our structures and functional recordings show that GABA is a high-efficacy ligand, activating over 80% of receptors without requirement of positive allosteric modulators. In accordance, etomidate binding to the α5β3-EM results in little change at a local or global scale, consistent with a conformational selection mechanism. It remains unclear whether this phenomenon represents a generalizable mechanism in the larger family, given that the synaptic α1β2γ2 subtype with GABA alone appears partially contracted at the 9′ activation gate relative to GABA-etomidate structures. Substituted cysteine accessibility measurements have indicated that general anesthetics stabilize a unique conformational state of α1β2γ2 versus GABA alone^32^, suggesting subtype-specific differences in potentiation.

The resting structure of α5β3-EM we obtained in the absence of fiducials provides a framework for structural characterization of heteromeric GABA_A_Rs which lack established antibodies or nanobodies suitable for use in cryo-EM. Interestingly, though purified in the absence of divalent ions, the resting state we obtained displayed clear density attributable to Zn^2+^ at the 17**′** of M2. The Zn^2+^ block of αβ-type GABA_A_Rs is well known and is a distinguishing feature of αβ-type compared to αβγ-type GABA_A_Rs and plays a role in pathological conditions associated with increased excitability. Loss of sensitivity in the αβγ-type GABA_A_Rs likely comes from the replacement of a β subunit with the γ subunit which lacks the 17**′** histidine. The structure and stoichiometry of α5β3γ-type receptors is still unknown, though the resistance to Zn^2+^ block and high cooperativity in GABA response hint at a ratio of 2:2:1 α5β3γ similar to the synaptic α1βXγ2 receptors. Further work on α5β3γ-type receptors is necessary to confidently define the assembly stoichiometry.

The presence of HEPES in the β-β interface has also been reported in previous structural studies, though its functional relevance is not known. HEPES could be modeled in both the resting and GABA bound states of the receptor though the binding pocket undergoes substantial rearrangement upon binding of GABA and Mb25. However, as we were unable to obtain the GABA-bound structure in the absence of Mb25, it is unclear how much of the conformational change at the β-β interface can be attributed to GABA alone. Given that it has been demonstrated to bind histamine, and activate both β3 homopentamers and α4β3δ GABA_A_Rs, the β-β interface may represent an intriguing new avenue for selectively targeting extrasynaptic GABA_A_Rs.

The density in the neurotransmitter binding pocket we find in the resting state is consistent with previous studies which report similar serendipitous density and tentatively assigned it to butyrate. We demonstrate that butyrate does indeed oppose activation of the receptor by GABA and have thus modeled butyrate into this density. Butyrate only lacks the primary amine group of GABA so it is not surprising that it can bind in this pocket, however it is roughly 4 times smaller than traditional inhibitors like bicuculline. The wide range in size in molecules that bind to this pocket indicate a high degree of flexibility in loop C at rest that is consistent with the reduced resolution and decreased local scale from OccuPy analysis we observed at this site in the absence of GABA. Future drug design efforts may leverage this dynamic orthosteric site using computational approaches like ensemble docking or AlphaFold 3 to improve specificity in competitive antagonists. Together with our observations of subunit assembly, subtype specificity, and modulatory mechanisms, this work thus offers a basis for biophysical insights and pharmacological targeting in the diverse and critical GABA_A_Rs.

## METHODS

### Construct design for cryo-EM

We designed α5-EM and β3-EM GABA_A_R constructs with numerous modifications to improve biochemical tractability. The α5-EM construct has the signal sequence from human α1 GABA_A_R followed in sequence by superfolder green fluorescent protein^33^, a Twin-Strep tag, and a recognition site for tobacco etch virus for expression and purification purposes^34^. Following the tags, the ECD+M1-M3 helices (residues 33-346) were separated from the M4 region (residues 421-462) of human α5 (UniprotID: P31644) by a linker sequence (SQPARAA). The β3-EM construct retained the sequence for human β3 (UniprotID: P28472) ECD+M1-M3 helices (residues 1-332), a linker containing the blue fluorescent protein mKalama1 (SQPGRA-mKalama1-GRAA), the β3 M4 sequence (residues 447-473) followed by a two residue linker (SR), TEV protease site, and 6 Histidine tag.

### Expression in oocytes and electrophysiology

To increase expression in *Xenopus* oocytes, α5-EM and β3-EM were transferred into the pUNIV expression vector. Plasmids encoding α5-WT, β3-WT, α5-EM, or β3-EM GABA_A_R constructs were linearized using KpnI, and RNA was produced by *in vitro* transcription with a mMessage mMachine T7 Ultra transcription kit (Ambion) according to the manufacturer protocol including the tailing reaction. Stage IV oocytes from *Xenopus laevis* frogs (Ecocyte Bioscience) were coinjected with 0.5 ng (WT) or 10 ng (EM) of RNA for each subunit. Injected oocytes were incubated for 1-2 days at 13°C in post-injection solution (88 mM NaCl, 10 mM HEPES, 2.4 mM NaHCO_3_, 1 mM KCl, 0.91 mM CaCl_2_, 0.82 mM MgSO_4_, 0.33 mM Ca(NO_3_)_2_, 2 mM sodium pyruvate, 0.5 mM theophylline, 0.1 mM gentamicin, 17 mM streptomycin, 10,000 u/L penicillin, pH 7.5) before use in two-electrode voltage clamp (TEVC) measurements.

For TEVC recordings, glass electrodes were pulled and filled with 3 M KCl to give a resistance of 0.5–1.5 MΩ and used to clamp the membrane potential of injected oocytes at −70 mV with an OC-725C voltage clamp (Warner Instruments). Oocytes were maintained under continuous perfusion with Ringer’s solution (123 mM NaCl, 10 mM HEPES, 2 mM KCl, 2 mM MgSO_4_, 2 mM CaCl_2_, pH 7.5) at a flow rate of 1.5 mL/min. Buffer exchange was performed using a gravity-fed, digitally-controlled solution exchange system (Scientific Instruments) and currents were digitized at a sampling rate of 2 kHz and lowpass filtered at 10 Hz with an Axon CNS 1440A Digidata system controlled by pCLAMP 10 (Molecular Devices). Concentration-response curves were fitted by nonlinear regression to the Boltzmann equation with variable slope using Prism 9.4 (GraphPad Software). Each reported value represents the mean and the standard error of the mean for three to four oocytes.

### Baculovirus production

Baculovirus was produced as previously described with several modifications. Briefly, chemically competent DH10BacVSV cells (Geneva Biotech) were transformed with a modified pEZT-BM vector encoding either the α5-EM or β3-EM GABA_A_R subunit and incubated at 37° C on transposition plates until the distinction between blue and white colonies was apparent^35^. White colonies for each subunit were picked and grown in suspension overnight and bacmid DNA was isolated as described previously^36^. Bacmids encoding α5-EM or β3-EM GABA_A_Rs were used to transfect TriEX Sf9 cells (Novagen) in 6-well plates using Cellfectin II (Fisher) according to manufacturer instructions except that 3 to 5 μg of bacmid was used. Supernatant containing the P1 virus was harvested following 96 h incubation at 27C. Virus was amplified by infecting 30-mL suspension Sf9 cell cultures^36^. Due to the slow amplification of the virus in this step, it was necessary to continue splitting cells to maintain cell density between 1 and 4 × 10^6^ cells/mL until ∼1% of cells showed red fluorescence due to the constitutive mCherry in the bacmid. Following this, cells were allowed to grow until the cultures were visibly red. P2 virus was further amplified by infecting 100 mL cultures of Sf9 cells to generate P3 virus that was used for transducing HEK cells. The structures obtained in this study were obtained from a total of three rounds of P3 virus production from the same P2 stocks.

### Expression and purification

Baculoviruses encoding the α5-EM or β3-EM subunits were mixed at a ratio of 1.2:1 prior to infection of Expi293F GnTl^-^ cells for protein production with 2.75% v/v virus-to-cell culture at a density of 3 × 10^6^ cells/mL. After 6 h incubation at 37°C, 5 mM sodium butyrate was added to cultures and flasks were moved to 30°C. Cells were harvested 48 h post-transduction and washed once with phosphate buffered saline before flash-freezing cell pellets in liquid nitrogen.

All steps of protein purification and grid preparation were performed at 4°C or on ice. Cell pellets from 2 L culture were resuspended in 100 mL cell resuspension buffer (300 mM NaCl, 40 mM HEPES pH 7.5 and 2 cOmplete protease inhibitor tablets (Roche)) and sonicated to lyse cells. Cell lysate was spun at 50,000 x g for 45 min to pellet crude membranes. The membrane pellet was resuspended in 100 mL 2X solubilization buffer (600 mM NaCl, 80 mM HEPES, pH 7.5 with 2 tablets of cOmplete protease inhibitor) and homogenized via sonication. Solubilization was initiated by the addition of 100 mL 2X detergent mixture (2% lauryl maltose neopentyl glycol (LMNG), 0.2% cholesteryl hemisuccinate (CHS)) and mixture was stirred for 2 h and 30 min. Solubilized membranes were clarified by centrifugation at 50,000 x g for 45 min. The supernatant was applied to 4 mL Streptactin Superflow resin (IBA) that was previously equilibrated in Buffer A (300 mM NaCl, 20 mM HEPES, 0.005% LMNG, 0.0005% CHS, pH 7.5) and batch-bound for 90 min with gentle mixing. The resin was washed with 20 column volumes of Buffer A and eluted using 5 column volumes Buffer A with 10 mM desthiobiotin (Sigma). The affinity-purified protein was concentrated and further purified by size-exclusion chromatography on a Superose 6 increase 10/300 column (Cytiva) with a mobile phase of 100 mM NaCl, 20 mM HEPES, 0.005% LMNG, 0.0005% CHS, pH 7.5. Peak fractions were pooled and concentrated to ∼3 mg/ml for grid preparation or nanodisc reconstitution.

### Nanodisc reconstitution

The SapA expression plasmid was a gift from Salipro Biotech AB. SapA was purified according to previously published protocols^37^. For the reconstitution of α5β3-EM into nanodiscs, purified α5β3-EM GABA_A_Rs, SapA and polar brain lipid (Avanti) were mixed at a molar ratio 1:15:150. The mixture was incubated on ice for 1 h before Bio-Beads SM-2 resin (Bio-Rad) was added and the mixture was gently shaken overnight at 4°C. After ∼16 hour incubation with biobeads, the supernatant containing SapA nanodiscs was collected and further purified by size-exclusion chromatography on a Superose 6 column with buffer containing 20 mM HEPES pH 7.5, 100 mM NaCl. Peak fractions were pooled and concentrated to ∼1 mg/mL.

### Megabody expression and purification

Megabody 25 was purified as described previously. Cultures (500 mL) were grown to an OD of 2 in TB and induced overnight with 1 mM IPTG at 25C. Cells were harvested and resuspended in 80 mL resuspension buffer (20% sucrose, 0.5 mg/mL lysozyme, 50 mM Tris, 1 mM EDTA, 150 mM NaCl, pH 8) and incubated at 4C for 30 minutes. Cell debries was pelleted at 8,000 x g for 25 minutes. The supernatant containing periplasmic proteins was loaded onto 2 mL bed volume of Ni-NTA resin pre-equilibrated in wash buffer (10 mM Tris, 140 mM NaCl, 5 mM imidazole, pH 7.3). Column was washed with 15 mL wash buffer before elution in wash buffer supplemented with imidizole to a final concentration of 500 mM. Affinity purified megabody was desalted by running a Superdex 200 size exclusion chromatography in 140 mM NaCl, 10 mM Tris, pH 7.3. Purified protein was concentrated to a concentration of 4 mg/mL and flash frozen until use in grid preparation.

### Grid preparation and EM data acquisition

For the resting state receptor without Mb25, fluorinated foscholine 8 (FFC-8) (Anatrace) was added to 2 mM before grid freezing. Samples prepared with Mb25 showed no preferred orientation even in the absence of FFC-8 and could be prepared at a lower protein concentration as noted above. Purified α5β3-EM GABA_A_R (3 μL) with the specified ligands was applied to a glow-discharged R1.2/1.3 400 mesh Au grid (Quantifoil), blotted for 2 s with force 0 and plunged into liquid ethane using a Vitrobot Mark IV (Thermo Fisher Scientific) at 22°C. Mb25 was added to samples at a final concentration of 0.4 mg/mL corresponding to a 3 fold molar excess above the concentration of the receptor. Mb25 and etomidate were added 45 minutes prior to freezing grids. GABA was added immediately before blotting corresponding to ∼10-15 seconds of exposure prior to freezing for most grids. One grid was prepared with GABA and Mb25 added together and incubated 45 minutes prior to freezing. Electron micrographs were collected at 300 kV using a Titan Krios (Thermo Fisher Scientific) electron microscope with a K3 Summit detector (Gatan) operating at 130,000x magnification corresponding to 0.648 or 0.65 Å/px using the EPU automated collection software (Thermo Fisher Scientific). Micrographs were recorded over a period of 1.1 to 1.4 s and fractioned into 40 frames for a total fluence of ∼40 to 65 e^-^/Å^2^ at the sample level as indicated in Table 1.

**Table 1.**
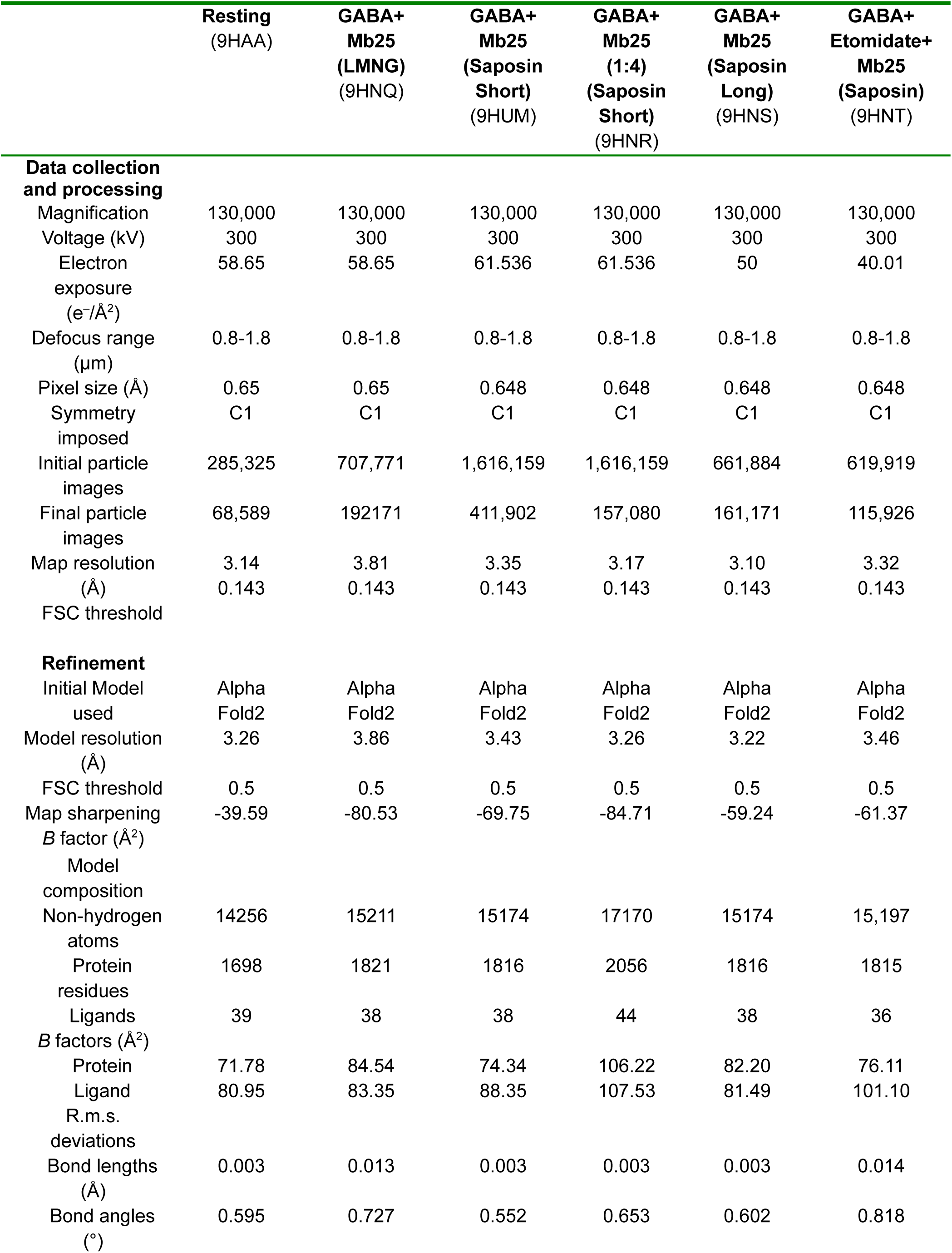
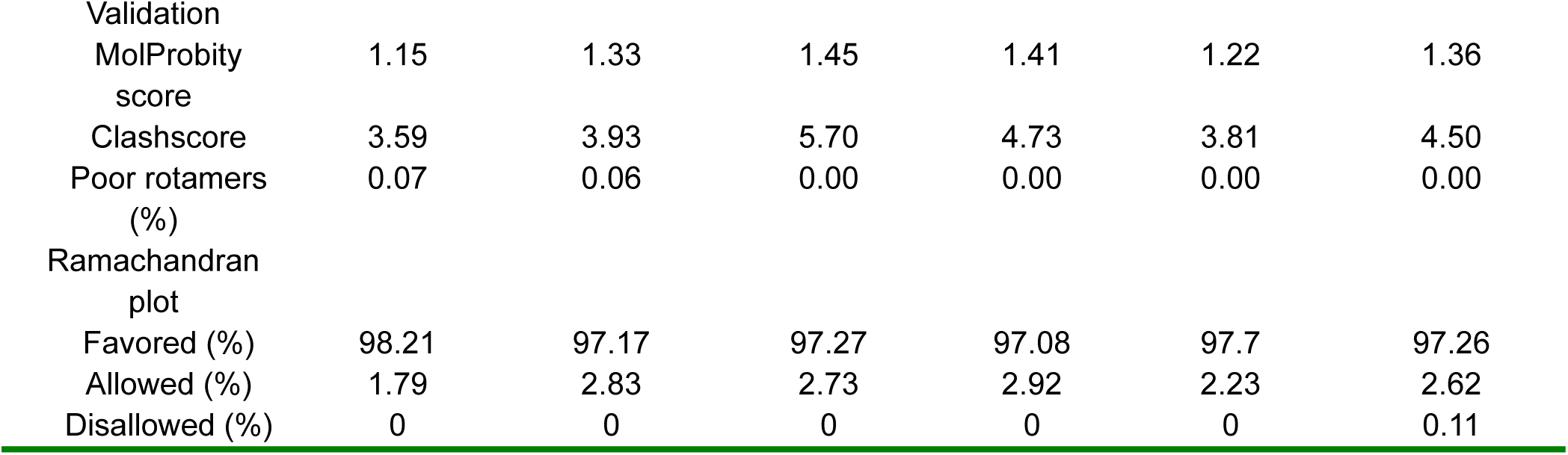
Cryo-EM data collection, refinement and validation statistics.

**Table 2:** Buried surface area for different interfaces in the resting and GABA-bound states in detergent micelles. All values are in Å^2^.

|  | Resting |  |  |  | GABA-bound |  |  |  |
| --- | --- | --- | --- | --- | --- | --- | --- | --- |
| Interface (chain) | ECD | TMD | ECD-TMD | FULL | ECD | TMD | TMD-ECD | FULL |
| $\beta$ (B)- $\alpha$ (A) | 1363.8 | 1288.5 | 94.0 | 2721.8 | 1573.5 | 1134.5 | 113.98 | 2780.4 |
| $\beta$ (C)- $\beta$ (B) | 1238.7 | 1372.6 | 115.2 | 2698.8 | 1426.9 | 1121.4 | 147.53 | 2651.1 |
| $\alpha$ (D)- $\beta$ (C) | 1535.2 | 1190.6 | 138.4 | 2796 | 1538.6 | 1122.6 | 122.37 | 2720.1 |
| $\beta$ (E)- $\alpha$ (D) | 1424.8 | 1342.5 | 106.5 | 2837.9 | 1555.6 | 986.4 | 106.61 | 2616.4 |
| $\alpha$ (A)- $\beta$ (E) | 1448.5 | 1188.1 | 125.0 | 2716.1 | 1519.6 | 1084.5 | 119.53 | 2683.7 |
| Total |  |  |  | 13770.6 |  |  |  | 13451.7 |

### Image processing

Dose-fractionated images in super-resolution mode were internally gain-normalized and binned by 2 in EPU during data collection. Beam-induced motion was corrected in Relion 4.0^38^. The contrast transfer functions (CTF) were estimated for motion-corrected micrographs using CTFFIND4.1^39^ and particles were automatically picked using the internal model in Topaz 0.2.5^40^. Picked particles were binned 4×4 and subjected to two rounds of 2D classification in Relion 4.0 on particles to remove empty nanodiscs and particles on carbon. Classes with recognizable channel features were re-extracted and centered by applying alignment offsets. These particles were used to generate initial models in CryoSPARC 4.0^41^. Further processing was done in Relion 5.0, including 3D classification without alignment to check the compositional and structural heterogeneity of classes. Multiple rounds of CtfRefine and polishing were executed to improve resolution before a final refinement using Blush regularization. For the resting state dataset, symmetry expansion was performed before the final refinement to improve alignment of the heteromeric complex as shown in Figure S3.

### Model building

Initial models of α5-EM and β3-EM monomers were created using AlphaFold2 The α5-EM or β3-EM sequences were modelled with ColabFold v1.2.0 for prediction of monomeric structures. Each subunit was fitted using Coot^42^ and manually edited to improve the local fit to density. Crystallographic information files (cif) for ligands were prepared from isomeric SMILES strings using Grade2^43^. Glycans were modeled based on previously assigned glycosylation patterns of α5 and β3, removing unresolvable moieties on the vestibular α5 glycan. Final models were optimized using real-space refinement in PHENIX^44^ and validated by MolProbity^45^. Subunit interfaces were analyzed using the PDBePISA server^46^, and pore radius profiles were calculated using the channel annotation package (CHAP) version 0.9.1 in Gromacs 2018. Hydrophobic estimates were calculated using MLPP^47^. Structure figures were prepared using UCSF ChimeraX^48^ and VMD.

## Resource Availability

### Lead Contact

Further information and requests for resources and reagents should be directed to and will be fulfilled by the lead contact, Erik Lindahl.

### Materials Availability

All unique/stable reagents generated in this study are available from the lead contact without restriction.

### Data and Code Availability

Cryo-EM density maps have been deposited in the Electron Microscopy Data Bank under accession numbers EMD-51980 (Resting state), EMD-52312 (GABA-bound in LMNG), EMD-52416 (GABA-bound in nanodisc, short application), EMD-52313 (GABA-bound in nanodiscs, 1 to 4 stoichiometry), EMD-52314 (GABA-bound in nanodisc, long application), EMD-52315 (GABA- and etomidate-bound in nanodisc). Model coordinates have been deposited in the Protein Data Bank under accession numbers PDB 9HAA (Resting state), 9HNQ (GABA-bound in LMNG), 9HUM (GABA-bound in nanodisc, short application), 9HNR (GABA-bound in nanodiscs, 1 to 4 stoichiometry), 9HNS (GABA-bound in nanodisc, long application), 9HNT (GABA- and etomidate-bound in nanodisc).

## Acknowledgements

We thank members of Molecular Biophysics Stockholm for feedback on the project and manuscript, and staff at the Swedish National Cryo-EM Facility for data collection and support. The data were collected at the Cryo-EM Swedish National Facility funded by the Knut and Alice Wallenberg, Family Erling Persson and Kempe Foundations, SciLifeLab and Stockholm University. JC was supported by an EMBO Postdoctoral Fellowship, CF by grant FV-5.1.2-0523-19 from Stockholm University, RJH, and EL by grants from the Knut and Alice Wallenberg Foundation (2023.0254), the Swedish Research Council (2019-02433, 2021-05806) and Swedish e-Science Research Center.

## Author Contributions

Conceptualisation: JC, RJH; methodology: JC, CF; validation: JC, CF; investigation: JC, CF; resources: JS; data curation: JC; writing - original draft: JC; writing - review and editing: JC, CF, RJH, EL; visualization: JC, RJH; supervision: RJH, EL; project administration: JC, RJH, EL; funding acquisition: JC, EL.

## Inclusion and Diversity

One or more of the authors of this paper self-identifies as a gender minority in their field of research. We support inclusive, diverse, and equitable conduct of research.

