## Extended Data Figures for "Structural basis for activation and potentiation in a human α5β3 GABA_A_ receptor"

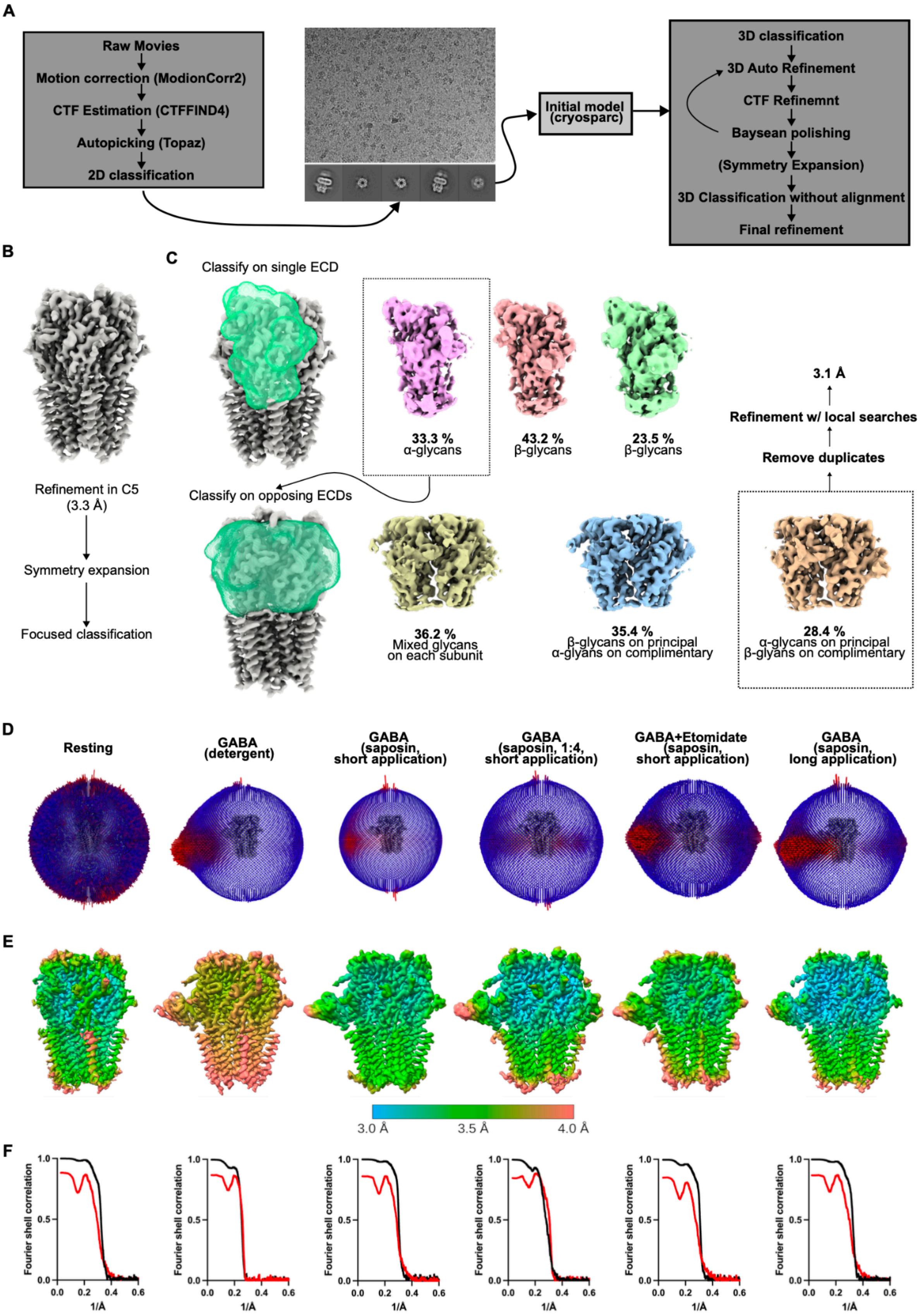
A
Raw Movies
Motion correction (ModionCorr2)
CTF Estimation (CTFFIND4)
Autopicking (Topaz)
2D classification
Initial model (cryosparc)
3D classification
3D Auto Refinement
CTF Refinemnt
Baysean polishing
(Symmetry Expansion)
3D Classification without alignment
Final refinement
B
Refinement in C5 (3.3 Å)
Symmetry expansion
Focused classification
C
Classify on single ECD
33.3 % α-glycans
43.2 % β-glycans
23.5 % β-glycans
Classify on opposing ECDs
36.2 % Mixed glycans on each subunit
35.4 % β-glycans on principal α-glyans on complimentary
28.4 % α-glycans on principal β-glyans on complimentary
Remove duplicates
Refinement w/ local searches
3.1 Å
D
Resting
GABA (detergent)
GABA (saposin, short application)
GABA (saposin, 1:4, short application)
GABA+Etomidate (saposin, short application)
GABA (saposin, long application)
E
3.0 Å
3.5 Å
4.0 Å
F
Fourier shell correlation
1/Å

**Figure S1. CryoEM processing workflow**

(A) Data processing workflow for datasets in this study

(B) Symmetry expansion workflow for reconstruction of heteromeric receptor without fiducial

(C) Classification strategy used in symmetry reconstruction first classifying on a single subunit to identify particles with α5-specific glycans at a given ECD position. This was followed by a second round of classification on the opposing two ECDs to isolate particles with correct global alignment. This analysis resolved particles with two clear α5 subunits and we were unable to isolate a class with only a single α5 subunit, indicating that the dominant stoichiometry is 2 α5 to 3 β3. Following the classification steps, a final refinement with local searches was performed to reconstruct the full density map.

(D) Angular distribution maps for the final reconstructions used in this study.

(E) Local resolution of electron density maps for each reconstruction.

(F) Map-to-map (black) and model-to-map FSC curves for each reconstruction

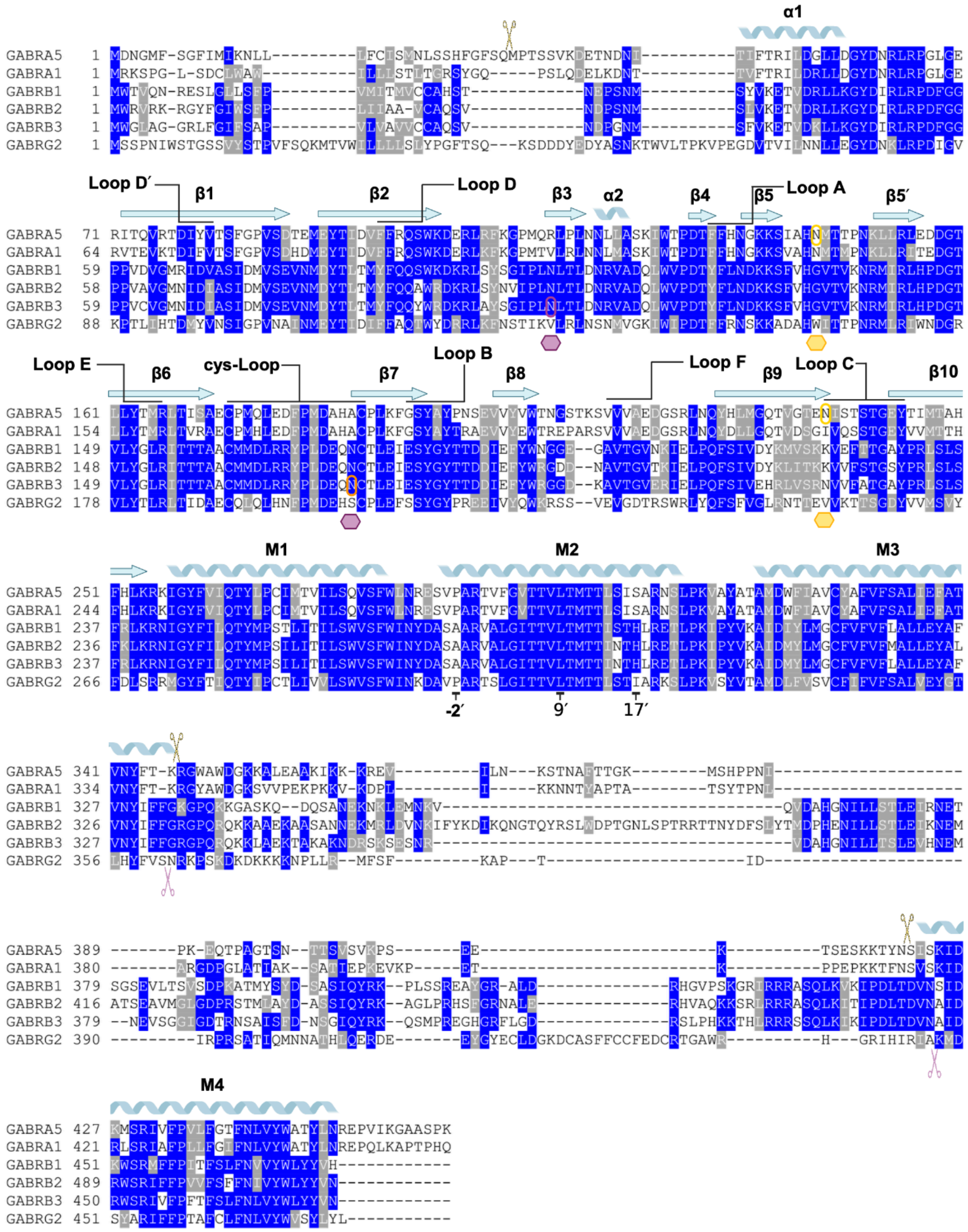

**Extended Data Figure 2. Sequence alignment of relevant GABAAR subunits**

Sequence alignment for α5 and β3 subunits with other GABAAR subunits relevant in this study. Glycans specific to α5 are marked with yellow hexagons while glycans specific to β3 are marked with purple hexagons. Scissors above the sequence indicates where α5 sequence was modified while scissors below sequence indicates modification positions of β3.

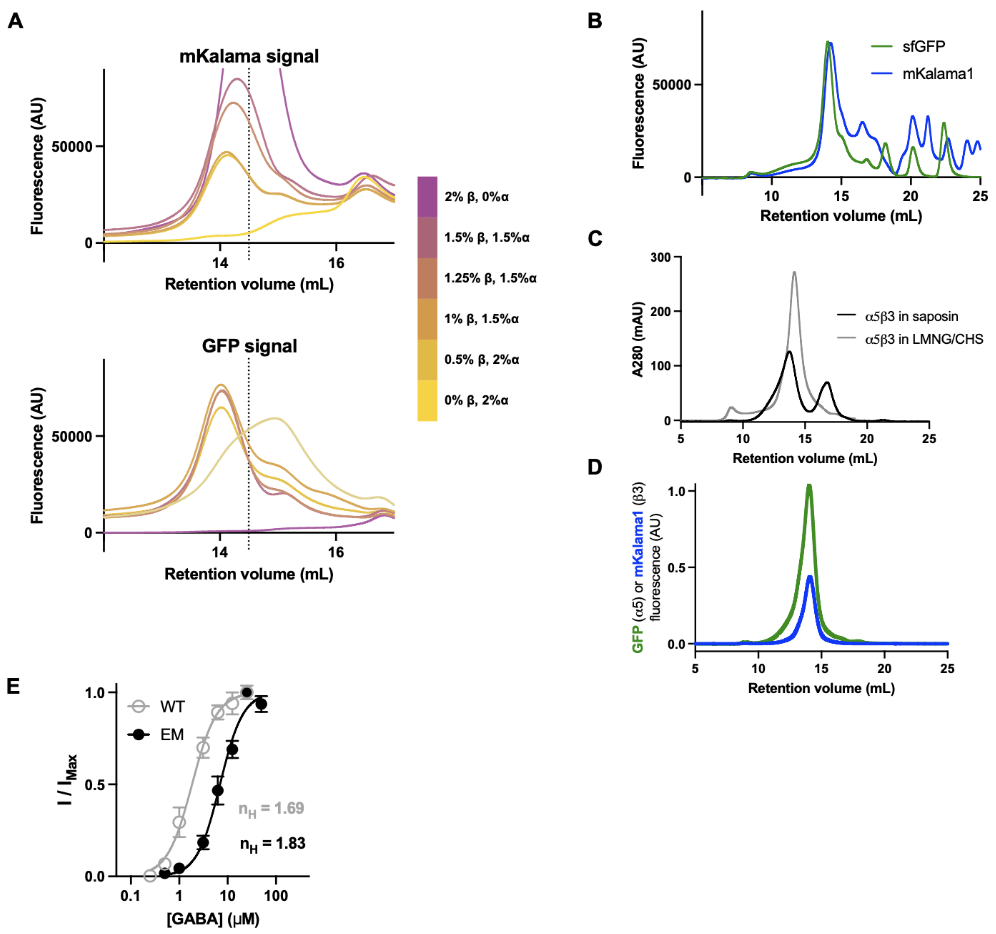


**Extended Data Figure 3. Biochemical and functional characterization of α5β3 GABAARs**

(A) Titration of viruses used for transduction. Suspension cultures were transduced with different ratios of α5-EM and β3-EM viruses and cells were assessed with FSEC to determine optimal ratio of viruses to use in large scale cultrues. The upper plot shows the FSEC traces in the blue detector (nm excitation, nm emission) and lower traces show the FSEC profiles in the green detector (480 nm excitation, 510 nm emission) to detect mKalama-labeled β3-EM and sfGFP-labeled α5-EM, respectively. Traces with the same color in the upper and lower plots come from the same sample as indicated in the color scale at the right.

(B) FSEC profile of the condition used for large scale expression of α5β3-EM with the blue curve representing the fluorescence signal in the blue detector and green curve coming from the green detector.

(C) SEC profile showing α5β3-EM purified in detergent (grey) or nanodiscs.

(D) FSEC profile of the cryoEM sample of α5β3-EM in nanodiscs showing coelution of GFP and mKalama1 indicating a heteromeric assembly.

(E) Concentration-response curves for α5β3-EM (black) and α5β3-WT (grey) expressed in *Xenopus* oocytes.

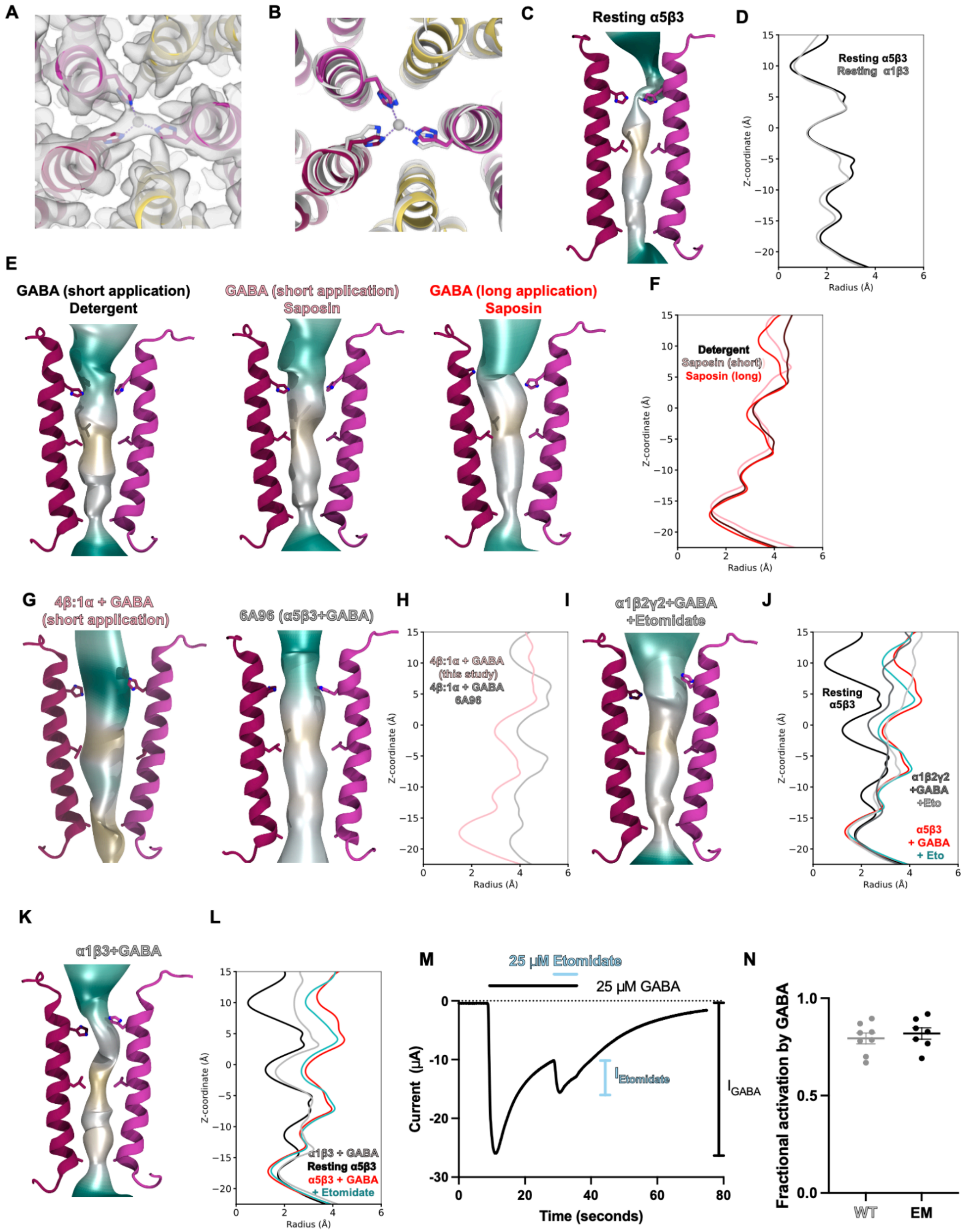


**Extended Data Figure 4. Comparison of pore properties of α5β3-EM with other receptors**

(A) Density map (grey transparent surface) overlaid with the atomic model for the resting state receptor showing a non-protein density in the pore consistent with bound $Zn^{2+}$

(B) Alignment of the pores of α5β3-EM (yellow/magenta) and α1β3 (grey) in resting states showing the side chains of H267.
(C) CHAP pore profile of the resting α5β3-EM structure colored by hydrophobicity with cyan representing more polar and yellow more hydrophobic. Side chains are shown for A248, L259, and H267 for opposing β3 subunits.
(D) CHAP pore profile as a function of z-coordinate relative to L259 for the resting state of α5β3-EM (black) or α1β3 (grey)
(E) Structural comparison of CHAP pore profiles for three GABA-bound α5β3-EM models obtained in detergent (left) or saposin nanodisc with short (middle) or long (right) applications of GABA prior to grid freezing.
(F) CHAP pore profile as a function of z-coordinate relative to L259 for the structures in E colored according to the labels.
(G) Structural comparison of CHAP pore profiles for GABA-bound α5β3-EM in a 1:4 stoichiometry from this study (left) compared with the previously published model (right).
(H) CHAP pore profile as a function of z-coordinate relative to L259 for the structures in G colored according to the labels.
(I) Structural overlay of CHAP pore profile with the GABA-bound α1β2γ2 structure with GABA and etomidate bound.
(J) CHAP pore profile as a function of z-coordinate relative to L259 for the structures of α5β3-EM with (red) or without (black) GABA in detergent compared to the GABA-bound α1β2γ2 structure with (light grey) or without (dark grey) etomidate.
(K) Structural overlay of CHAP pore profile with the GABA-bound α1β3 structure.
(M) Example trace showing estimation of the fraction of receptors directly activated by GABA. Fractional activation is defined as $I_{GABA}/(I_{GABA}+I_{Etomidate})$
(N) Fractional activation of α5β3-EM (black) and α5β3-WT (grey) by GABA as estimated using the method described in M. Solid bars indicate mean values with error bars showing standard error of 8 independent oocytes shown as individual points.

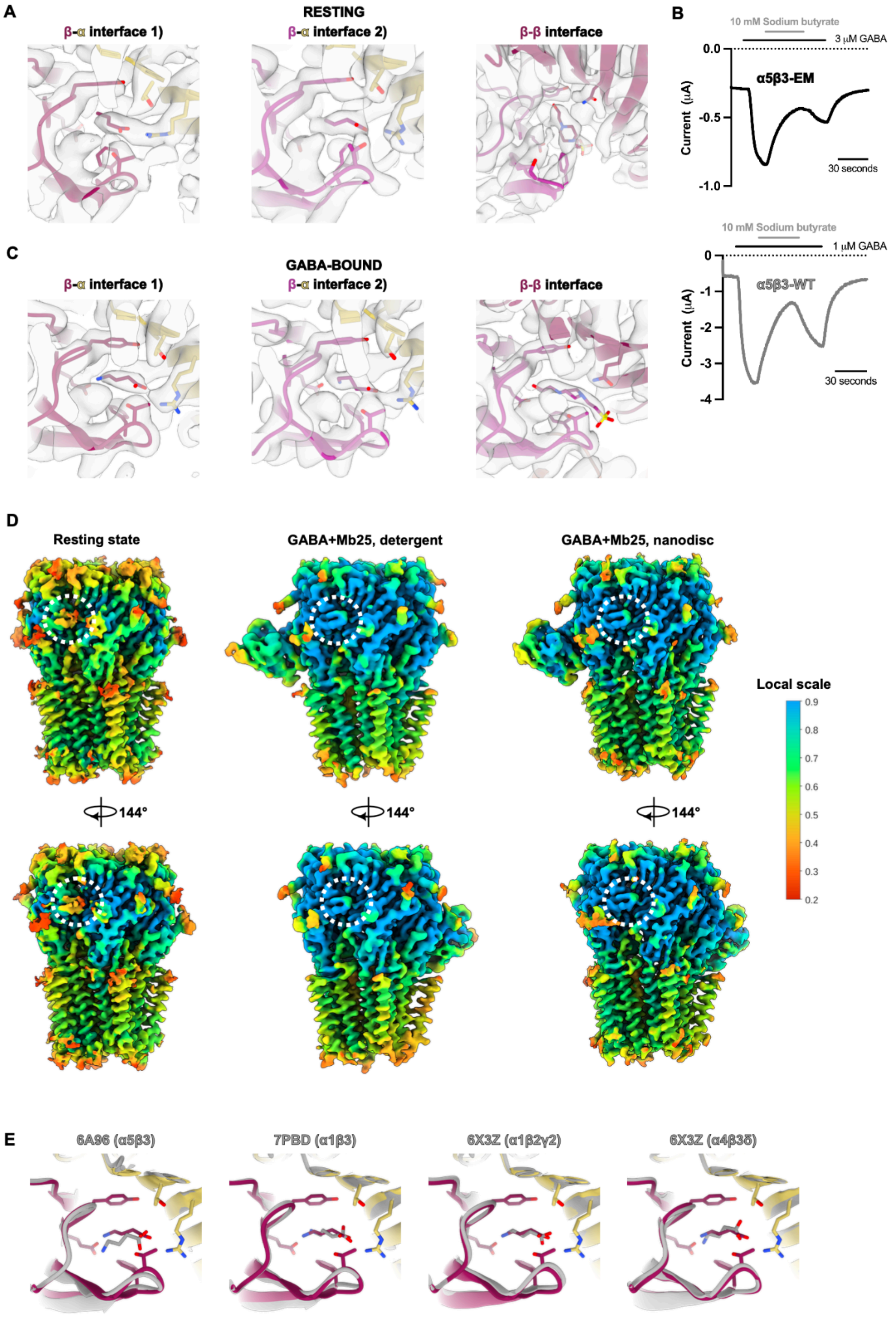


**Extended Data Figure 5. Comparison of neurotransmitter binding pocket across structures**
(A) Density map (grey transparent surface) overlaid with the atomic model for the resting state receptor showing a non-protein density in the β-α and β-β interfaces assigned to butyrate and HEPES, respectively. (B) Example two-electrode voltage clamp traces for α5β3-EM (top) and α5β3-WT (bottom) in response

to GABA application at ~EC50 with a coapplication of 10 mM sodium butyrate in the middle.
(C) Density map (grey transparent surface) overlaid with the atomi model for the GABA-bound state receptor showing a non-protein density in the β-α and β-β interfaces assigned to GABA and HEPES, respectively.
(D) OccyPy analysis of density maps for the resting state, and GABA-bound receptors in detergent (middle) or saposin nanodisc (right). Maps are colored according to local scale based on the scale bar on the left and the region corresponding to the neurotransmitter binding pocket is circled. Maps are rotated 144 degrees in lower panels to show second neurotransmitter site.
(E) Structural GABA binding pose for the α5β3-EM and previously resolved heteromeric GABA receptors.

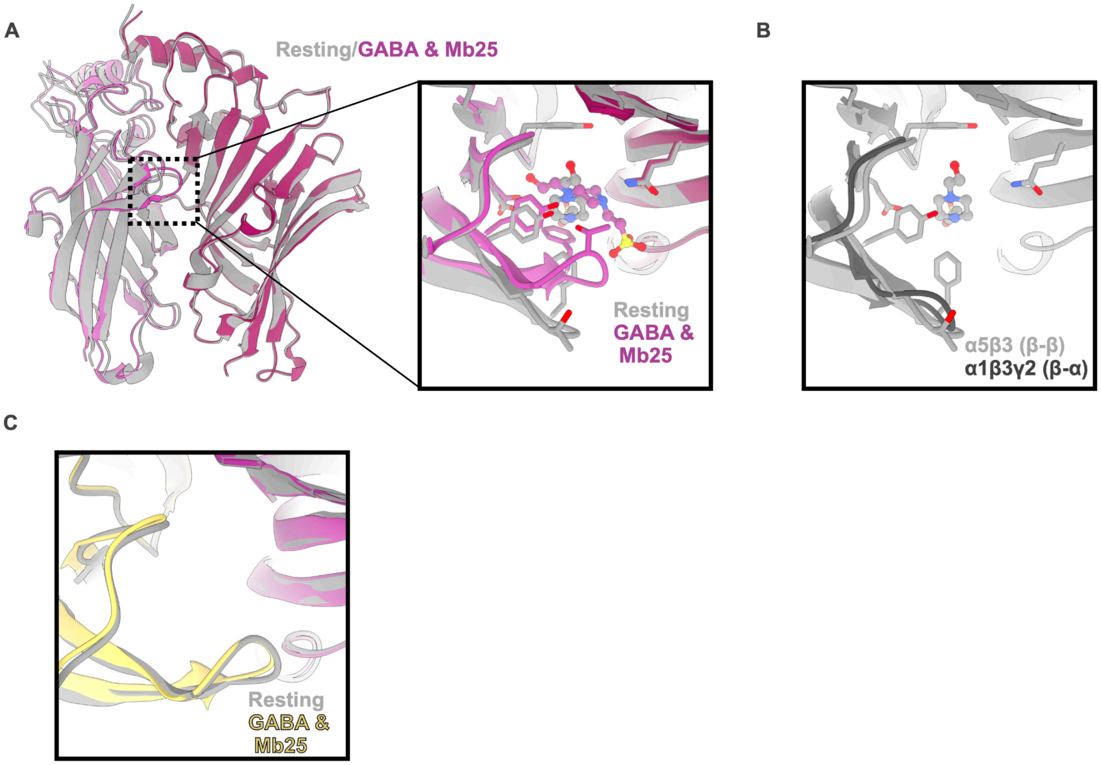


**Extended Data Figure 6. Rearrangements at other ECD interfaces**

(A) Left: ECDs of the β-β interface in the resting (light gray) and desensitized (light/dark magenta) α5β3-EM structures, aligned on the complementary subunits. Right: Zoomed view of the binding pocket occupied by HEPES in both states with relevant residues in the pocket interacting with the ligands are shown as sticks.

(B) Comparison of the β-β interface in the resting α5β3-EM structure (light gray) with the β-α interface of the α1β3γ2+PTX structure (dark gray)

(C) Comparison of the α-β interface in the resting α5β3-EM structure (light gray) with the desensitized (light/dark magenta) α5β3-EM structures

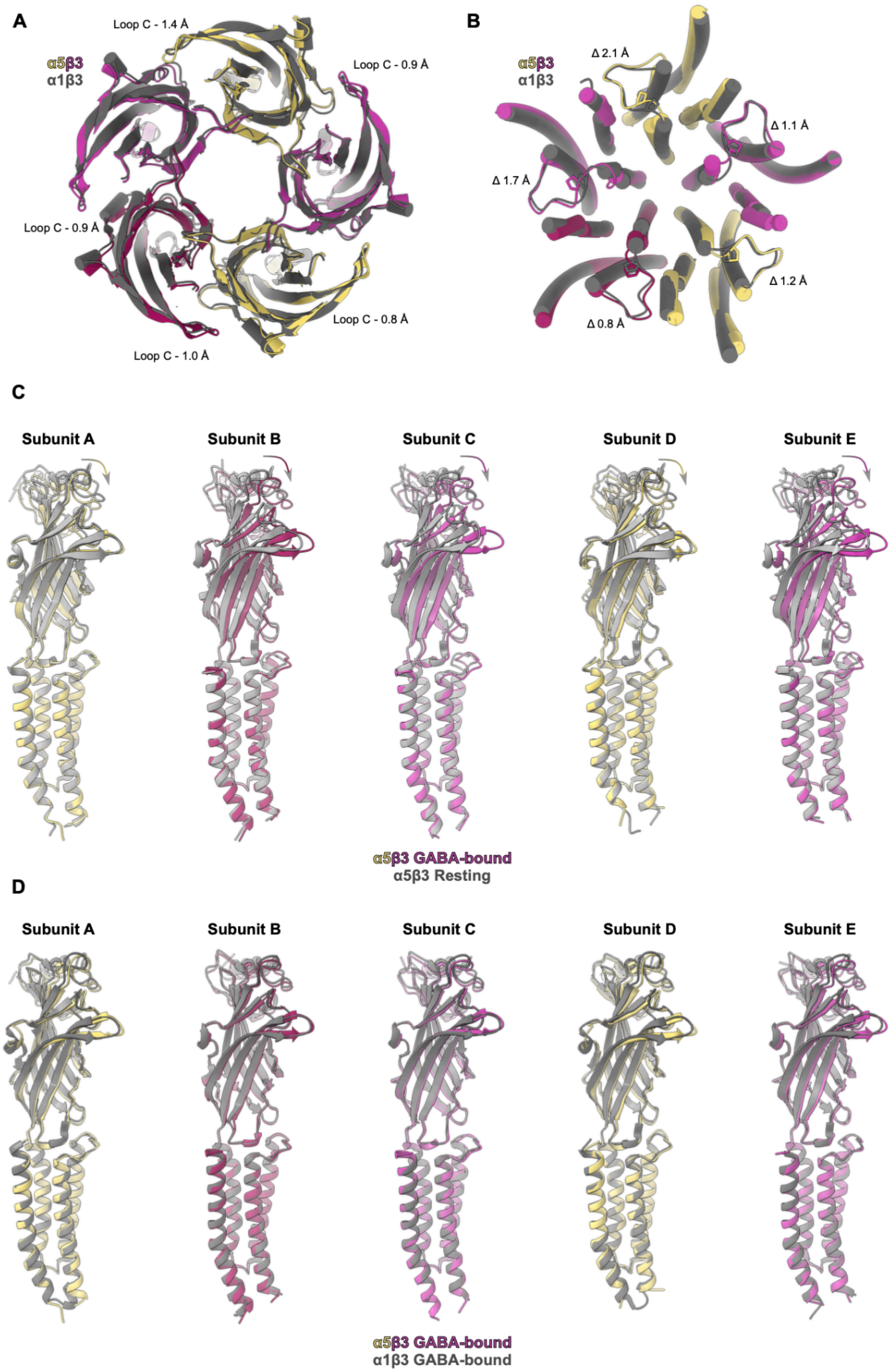


**Extended Data Figure 7. Comparison of conformations of GABA-bound α5β3-EM and α1β3**

(A) Overlay of ECDs from GABA-bound α1β3 (gray) and α5β3-EM (magenta/yellow) receptors aligned by their TMDs. The slice of the receptor displayed is the same as in Figure 4A. Distances measure the distances between CA atoms of the threonine at the tip of loop C.

(B) Overlay of TMD from GABA-bound α1β3 (gray) and α5β3-EM (magenta/yellow) receptors aligned by their TMDs. The slice of the receptor displayed is the same as in Figure 4C. Distances measure the distance between P281(α5)/P273(β3) in α1β3 compared to α5β3-EM.
(C) Overlay of individual subunits from resting (light gray) and GABA-bound (magenta/yellow) receptors aligned by their respective TMDs.
(D) Overlay of individual subunits from GABA-bound α1β3 (dark gray) and α5β3-EM (magenta/yellow) receptors aligned by their respective TMDs.

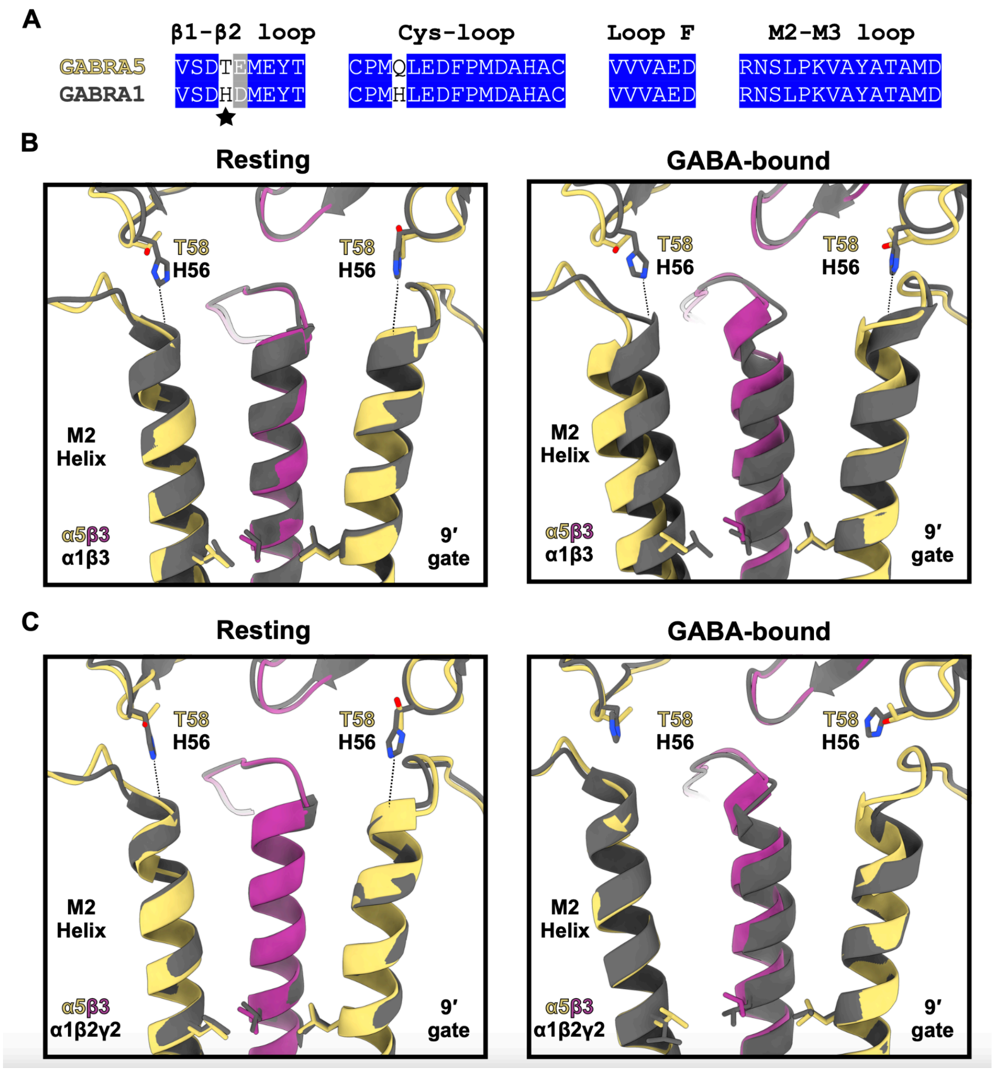


**Extended Data Figure 8. Differences in the coupling interface in α5 and α1 subunit-containing receptors**

(A) Sequence comparison for the critical regions at the ECD-TMD interface between α5 and α1 with the position of T58/H56 marked by a star.

(B) Overlay of M2 helices and β1-β2 loops for α1β3 (dark gray) and α5β3-EM (magenta/yellow) aligned by the regions shown. Left panel shows the comparison of the resting state structures and right panel shows comparison of GABA-bound structures. Dashed lines indicate distances between the side chain of H56 and backbone atoms of the M2 helix. Two subunits and all other residues of the TMD and ECD are hidden for clarity.

(C) Overlay of M2 helices and β1-β2 loops for α1β2γ2 (dark gray) and α5β3-EM (magenta/yellow) aligned by the regions shown. Left panel shows the comparison of the resting state structures and right panel shows comparison of GABA-bound α5β3-EM with GABA+etomidate-bound α1β2γ2. Dashed lines indicate distances between the side chain of H56 and backbone atoms of the M2 helix that are notably absent in the GABA+etomidate-bound structure. Two subunits and all other residues of the TMD and ECD are hidden for clarity.

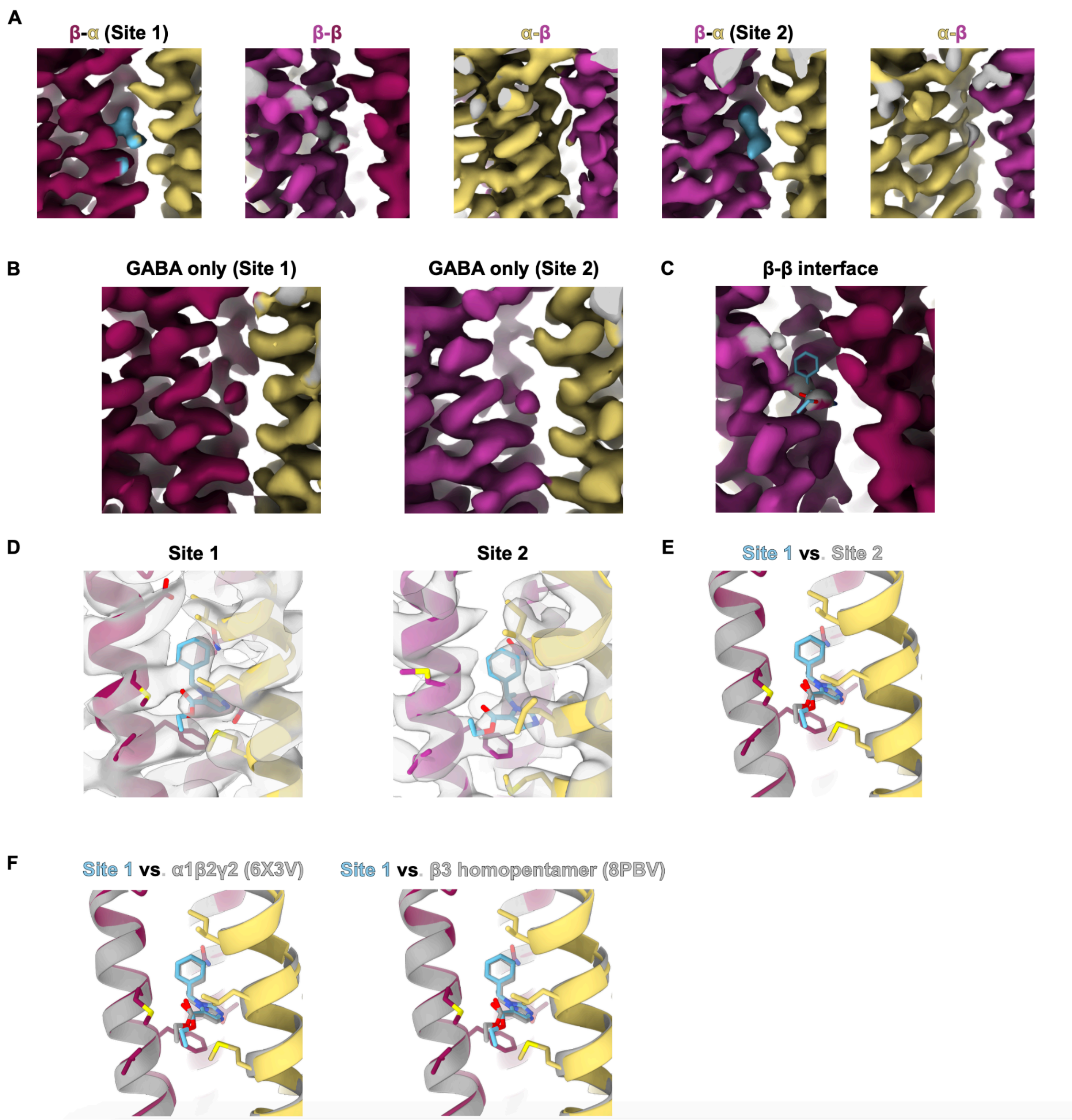


**Extended Data Figure 9. Etomidate binding pocket**

(A) Density map for the principal M3/complimentary M1 interfaces in the etomidate-bound structure colored according to the same scheme as Figure 1.

(B) Density map for the principal M3/complimentary M1 interfaces in the GABA-only structure colored according to the same scheme as Figure 1 shown only for the two β-α interfaces.

(C) Density maps for the principal M3/complimentary M1 β-β interface in the etomidate-bound structure colored according to the same scheme as Figure 1 overlaid with an etomidate model (cyan sticks) showing poor fit quality.

(D) Density map (grey transparent surface) overlaid with the atomic model for the etomidate-bound receptor showing the fit of etomidate (cyan sticks) in the density map.

(E) Overlay of the two etomidate binding sites at the β-α interfaces within the etomidate-bound α5β3-EM structure

(F) Overlay of the etomidate binding sites in α5β3-EM structure (site 1, magenta/yellow/cyan model) with the etomidate-bound α1β2γ2 (gray model, left) and β3 homopentamer (gray model, right).
